# Senolytic targeting of CAF-induced KRT17^+^ colon cancer cells inhibits metastatic invasion

**DOI:** 10.64898/2026.03.03.708913

**Authors:** Daisuke Shiokawa, Hiroaki Sakai, Yusuke Kanda, Hirokazu Ohata, Yusuke Asada, Hiroki Ochiai, Takeo Fukagawa, Shigeki Sekine, Yasushi Yatabe, Masato Morikawa, Hitoshi Nakagama, Koji Okamoto

## Abstract

Metastasis remains the leading cause of cancer mortality, yet effective strategies to eliminate metastasis-initiating cells are lacking. Here, we dissected intratumoral heterogeneity in colon cancer using single-cell analyses and identified a *KRT17^+^* slow-cycling cancer cell population that exhibits features of metastasis-initiating cells (*L1CAM*) and senescent-like cells (*CDKN2A, BCL2L1*). Spatial transcriptomics and immunostaining revealed that these cells localize at tumor-stroma interfaces, where they are closely associated with TGF-β1-producing subset of cancer-associated fibroblasts (CAFs) and exhibit SMAD3 activation. Mechanistically, TGF-β1 induces KRT17 expression in patient-derived cancer cells, while co-culture with CAFs drives the emergence of KRT17 migratory cells in a KRT17-dependent manner. Functionally, genetic ablation of KRT17 or senolytic targeting of BCL2L1 suppresses peritumoral invasion and liver metastasis in xenograft models. Clinically, KRT17 cells co-localize with TGF-β1 CAFs, and their co-expression with L1CAM correlates with advanced disease stage. These findings support a model in which stromal TGF-β signaling promotes the emergence of a KRT17 invasive cancer cell state with senescence-associated features and suggest that senolytic strategies may represent a potential approach to limit metastatic progression in colon cancer.

## INTRODUCTION

Metastasis to distant organs is the leading cause of cancer-related mortality and involves a multistep processes including local invasion, intravasation, survival in circulation, extravasation, and colonization at secondary sites (*1*). While each step requires distinct biological capabilities, the early emergence of cancer cells with metastatic potential— termed metastasis-initiating cells (MICs)—is considered a crucial determinant of metastatic progression (*1, 2*). In colorectal cancer, several molecular markers associated with MIC-like properties have been proposed (*3, 4*), among which L1 cell adhesion molecule (L1CAM) has been implicated in both metastatic dissemination and chemoresistance (*3*). However, biological characteristics, origin, and regulatory mechanism underlying MIC formation remain poorly understood.

Transforming growth factor-β (TGF-β) is a key regulator of tumor progression with context-dependent roles in metastasis (*5*). In early stages, TGF-β promotes epithelial-mesenchymal transition (EMT), thereby enhancing phenotypic plasticity and invasive capacity (*6, 7*). Additionally, TGF-β shapes the tumor microenvironment by activating fibroblasts into cancer-associated fibroblasts (CAFs), which further support tumor progression (*5*). These pleiotropic effects suggest that TGF-β signaling may contribute to epigenetic programming of cancer cells toward MIC states. Nevertheless, whether and how stromal-derived signals, particularly from CAFs, directly induce MIC formation in colorectal cancer remain unclear.

CAFs themselves comprise heterogeneous subpopulations, including myofibroblastic CAFs (myCAFs), inflammatory CAFs (iCAFs) (*8–11*), and antigen-presenting CAFs (apCAFs) (*12*). Although specific CAF subsets such as iCAFs have been linked to chemoresistance in rectal cancer (*13*), their role in promoting metastatic competence is not fully defined. Moreover, the tumor microenvironment involves complex interactions with immune cells beyond CAFs (*14*); for example, SPP^+^ macrophages cooperate with FAP^+^ fibroblasts to form desmoplastic niches (*15*), and metabolically-activated M2-like macrophages facilitate liver metastasis (*16*). These findings highlight the importance of cellular networks in shaping metastatic potential, yet their impact on MIC generation remains to be elucidated.

MICs share key properties with cancer stem cells (CSCs), including phenotypic plasticity, migratory capacity, and resistance to chemotherapy. CSCs can dynamically transition between functional states in response to environmental cues through epigenetic mechanisms (*17–19*). This phenotypic resemblance suggests a potential overlap between MIC and CSC populations.

In our previous work, we identified distinct slow- and fast-cycling CSC populations in colon cancer xenografts, with the slow-cycling population exhibiting enhanced chemoresistance (*20*). This population was characterized by elevated expression of a slow-cycling CSC gene signature, including PROX1, MEX3A, ZNF503, and KRT17 (*20*). Notably, Mex3a has been reported as a marker of reserve stem-like cells with increased chemoresistance (*21*), supporting the functional relevance of this cell state.

In this study, we extended our investigation and identified a KRT17^+^ colon cancer cell subpopulation that co-expresses metastatic markers and slow-cycling CSC-associated genes, suggesting a hybrid state with both stem-like and metastatic properties. We further demonstrate that CAF-derived TGF-β drives the generation of this subpopulation. Importantly, these cells exhibit features of cell senescence, and targeting these cells by senolytic treatment or KRT17 inhibition significantly suppressed cancer invasion and metastasis. These findings provide mechanistic insight into generation of cancer cells with invasive/metastatic capability and highlight a previously unrecognized link between stromal signaling, cellular senescence, and metastatic competence.

## RESULTS

### A subpopulation of *KRT17^+^* slow-cycling CSCs localizes at the tumor-stromal boundary

Previously, we identified *LGR5^+^*/*PROX1^+^*slow-cycling cells as a chemoresistant CSC population in mouse xenograft tumors generated after transplantation of patient-derived colon cancer spheroids (*20*). To further examine cellular diversity of this slow-cycling CSC population consists of one or multiple subpopulations, we transplanted cancer spheroids (CRC-6) into NOD/Shi-scid, IL-2RγKO (NOG) mice, isolated *EPCAM^+^*epithelial populations, and performed single-cell RNA sequencing (scRNA-seq) using the Chromium platform (10x Genomics). After quality control and normalization utilizing Seurat (*22*), we obtained transcription profiles for 2019 cancer cells (median count, 44337 UMIs/cells). We identified nine subpopulations through unsupervised graph-based clustering, which we visualized in a Uniform Manifold Approximation and Projection (UMAP) plot (Fig. 1A).

**Fig. 1.**
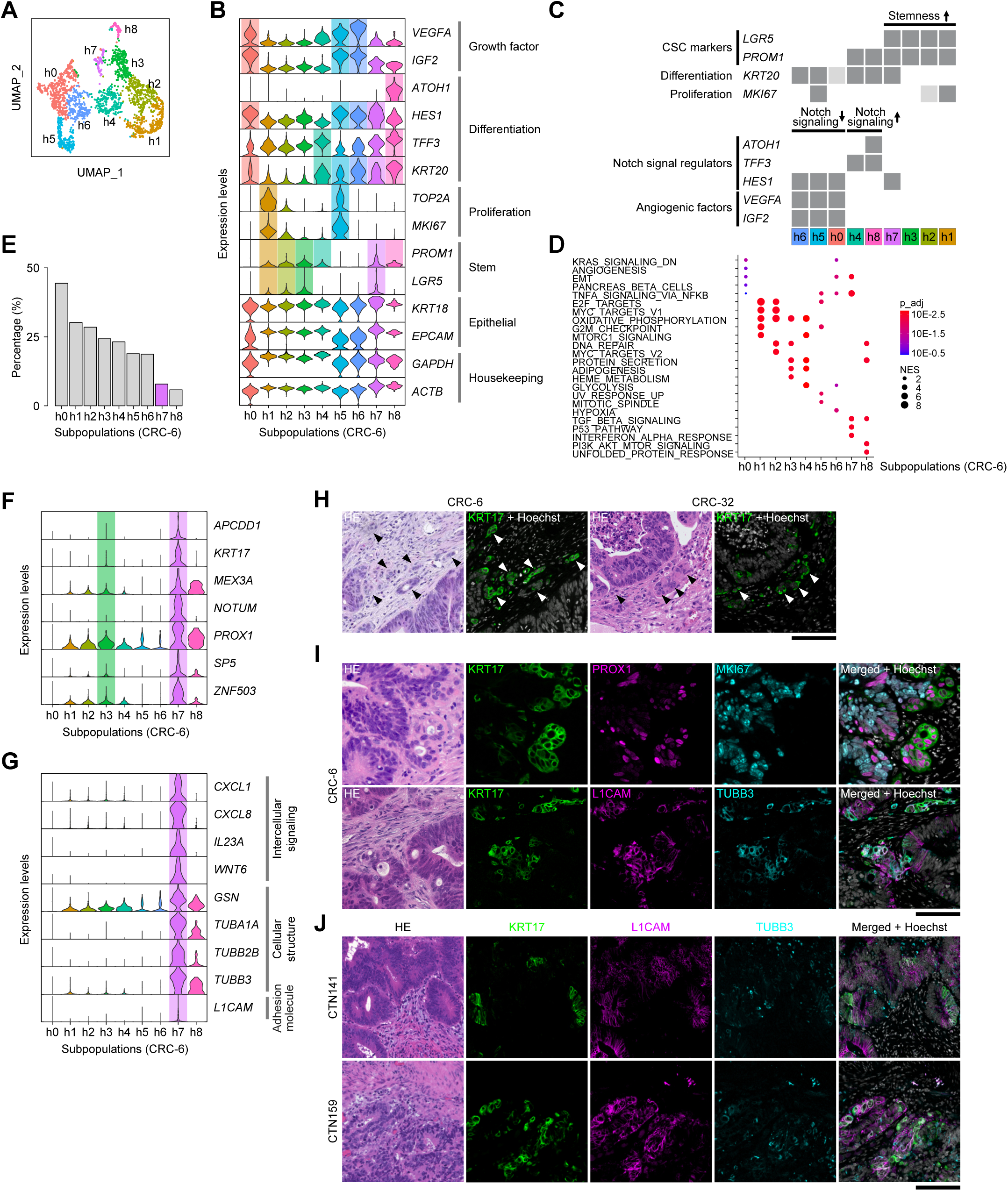
A subpopulation of *KRT17*^+^ slow-cycling CSCs localizes at the tumor-stromal boundary. (**A**) A UMAP plot of the *EPCAM*^+^ cancer cell subpopulations (h0–h8) in xenograft tumors (CRC-6). (**B**) Violin plots showing expression of representative cell type-specific marker genes in the cancer subpopulations. (**C**) Cell type annotation based on expression of the cell type-specific markers shown in B. (**D**) Gene Set Enrichment Analyses (GSEA) of the cancer subpopulations. The top five hallmark gene sets enriched in each subpopulation are shown. The adjusted *p*-value (p_adj) for each gene set is color-labeled. Dot size represents the normalized enrichment score (NES). (**E**) The relative proportion of each *EPCAM*^+^ cancer subpopulation is shown. The h7 subpopulation is highlighted in magenta. (**F**) Violin plots showing expression of slow-cycling CSC signature genes in the cancer subpopulations. (**G**) Violin plots showing expression of genes preferentially expressed in the h7 subpopulation. (**H**) H&E staining and KRT17 immunostaining of the xenograft tumors (CRC-6 and CRC-32). Arrowheads indicate KRT17^+^ invasive tumor cells. Scale bar: 100 μm. (**I**) H&E staining and immunostaining of the xenograft tumors (CRC-6) with the indicated antibodies. Scale bar: 100 μm. (**J**) H&E staining and immunostaining of surgical specimens of human colon cancers. Scale bar: 100 μm.

Next, we determined cellular identity of each subpopulation based upon the expression of specific marker genes (Fig. 1, B and C, and table S1). Four out of the nine subpopulations (h1-3, h7) showed detectable levels of *LGR5,* an established stem cell marker (*23*). One of these *LGR5^+^* populations (h1) expressed high levels of *MKI67* and *TOP2A* (Fig. 1B), as well as cell cycle-related signatures (Fig. 1D; E2F_targets, MYC_targets_V1), indicating that this population corresponded to actively cycling CSCs.

Among the other *LGR5^+^* populations, the h3 and h7 subpopulations are likely to correspond to the slow-cycling CSCs as they expressed low levels of cell cycle-related markers. Interestingly, the h7 subpopulation, which was smaller in number than the h3 subpopulation (Fig. 1E), was associated with an EMT signature (Fig. 1D). The slow-cycling *LGR5^+^* populations expressed all (h7) or some (h3) of the seven previously identified slow-cycling CSC signature genes (*20*) (Fig. 1F). Notably, genes expressed preferentially in the h7 subpopulation included *TUBB3* and *L1CAM* (Fig. 1G), markers of metastatic colon cancer cells (*3, 24, 25*), suggesting possible involvement of this subpopulation in metastasis.

To investigate whether the identified subpopulation expressing metastasis-associated genes was present in additional cases, we performed scRNA-seq of cancer cells from another patient-derived xenograft tumor (CRC-32) and obtained transcription profiles for 2179 cancer cells (median count, 20731 UMIs/cell). Clustering analyses identified nine cancer cell subpopulations (fig. S1A and table S2). The h2 subpopulation expressed *LGR5* together with high levels of *MKI67* and *TOP2A*, indicating that this subpopulation corresponded to actively cycling CSCs (fig. S1, B and C). Notably, the h7 subpopulation of CRC-32, which comprised a small proportion (fig. S1D), expressed some slow-cycling CSC signature genes (*KRT17*, *PROX1*, and *ZNF503*; fig. S1E), as well as the genes preferentially expressed in the h7 subpopulation of CRC-6 (Fig. 1G and fig. S1F), suggesting that the h7 subpopulation derived from CRC-6 and CRC-32 share similar expression profiles. Indeed, after integration of scRNA-seq data from CRC-6 and CRC-32 (fig. S1, G and H), the h7 subpopulation of CRC-6 and CRC-32 clustered into an identical population (c10 in the integrated data; fig. S1I). Thus, both xenograft tumors harbored the slow-cycling subpopulation expressing metastasis-related markers.

Next, to examine the spatial localization of the slow-cycling h7 subpopulation, we performed immunostaining of KRT17, which was uniquely expressed in the h7 subpopulation from both tumors (Fig. 1F and fig. S1E). KRT17 staining was observed near to the tumor-stromal boundaries, mainly at invasion fronts (Fig. 1H and fig. S1J). While KRT17 staining largely overlapped that of PROX1, a slow-cycling CSC signature marker, PROX1^+^ cells were also observed in glandular epithelia (Fig. 1I, upper panel). We did not observe MKI67 expression in KRT17^+^ cells (Fig. 1I, upper panel), which was consistent with the results of single-cell analyses (Fig. 1B).

As expected from the results of single-cell analyses, we observed extensive co-immunostaining of KRT17 with L1CAM and TUBB3 (Fig. 1I, lower panel). To examine expression of KRT17 and L1CAM in clinical specimens of human colon cancer, 12 surgical specimens of colon cancers (table S3) were subjected to immunostaining. We observed extensive co-localization of KRT17 with L1CAM and TUBB3 in six cases (Fig. 1J and fig. S1K). In other cases, only one of them was highly expressed (fig. S1L, lower 4 panels) or neither of them showed high levels of expression (fig. S1L, upper 2 panels). Thus, while expression levels of KRT17/L1CAM in clinical human cancers is variable, a significant fraction of clinical cancers co-express both proteins.

### Stratification of stromal populations within the colon tumors

To explore potential interaction of stromal cells with the identified metastasis-related cancer cells, we performed scRNA-seq analyses of non-tumor stromal cells from a xenograft tumor (CRC-6). We analyzed approximately equal numbers of FACS-sorted EpCAM^-^/CD45^+^ and EpCAM^-^/CD45^-^ cells (∼5×10^3^ cells each) using scRNA-seq (fig. S2A), and obtained transcription profiles for 719 EpCAM^-^/CD45^+^ cells (median counts, 18140 UMIs/cell) and 1195 EpCAM^-^/CD45^-^ cells (median counts, 2975 UMIs/cell). Subsequently we stratified them into six EpCAM^-^/CD45^-^ subpopulations (m0, m1, m3, m6, m8, and m9) and four EpCAM^-^/CD45^+^ subpopulations (m2, m4, m5, and m7) based upon their gene expression profiles (table S4), which were visualized them in a UMAP plot (Fig. 2A).

**Fig. 2.**
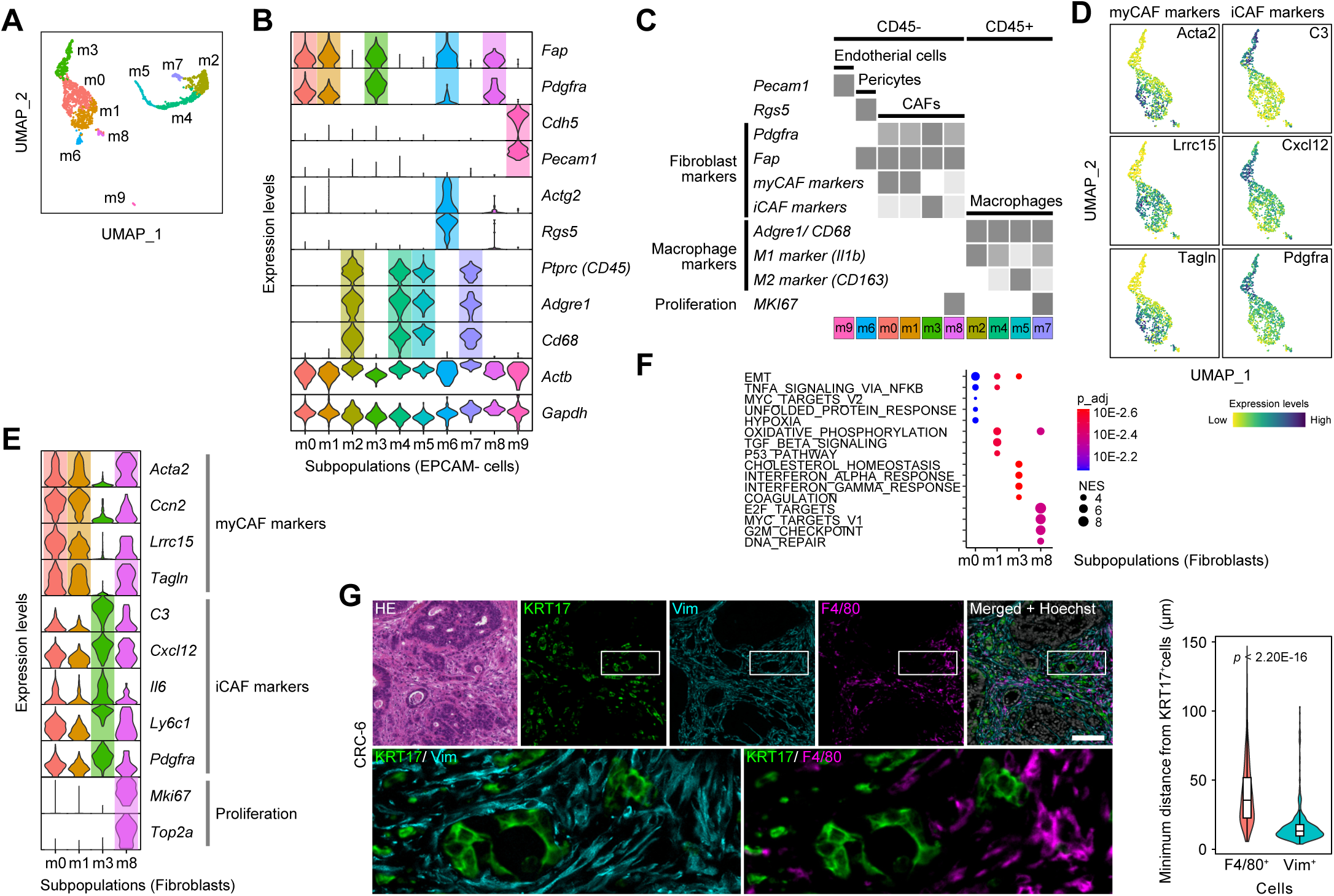
Stratification of stromal populations of the colon tumors. (**A**) A UMAP plot of the *EPCAM^-^* non-tumor subpopulations (m0–m9) in xenograft tumors (CRC-6). (**B**) Violin plots showing expression of representative cell type-specific marker genes in the non-tumor subpopulations. (**C**) Cell type annotation based on the expression of the cell type-specific markers shown in B. (**D**) UMAP Feature plots showing expression of the indicated marker genes for myCAF and iCAF in the fibroblast subpopulations (m0, m1, m3, and m8). (**E**) Violin plots showing expression of the indicated marker genes for myCAF, iCAF, and cell proliferation. (**F**) GSEA of the CAF subpopulations. the top five hallmark gene sets enriched in each subpopulation are shown. (**G**) H&E staining and immunostaining of the xenograft tumors (CRC-6). Magnified view of the White boxes in the upper panels is shown at the bottom. Scale bar: 100 μm. Violin plots of the minimum Euclidean distances from KRT17⁺ cells to F4/80⁺ and Vim⁺ cells are shown in the right panel.

The m6 and m9 subpopulations of the EpCAM^-^/CD45^-^ stromal cells likely represent pericytes and vascular endothelial cells, respectively, based upon their marker expression (*Rgs5* and *Pecam1*) (*26*) (Fig. 2, B and C). The other EpCAM^-^/CD45^-^ subpopulations (m0, m1, m3, and m8) expressed fibroblast markers *Fap* and *Pdgfra* (*27*) (Fig. 2, B and C), and have been categorized into several functional subtypes based upon expression of CAF markers (*8, 9, 11, 28*). Expression of specific markers indicated that the m0 and m1 populations represented myCAFs (Fig. 2, D and E). Elevated TGF-β signaling in the m1 subpopulation supports its identity as myCAFs (*29*) (Fig. 2F). Elevated expression of iCAF markers (*12*) (Fig. 2, D and E), as well as interferon responses (Fig. 2F), indicated that the m3 subpopulation represents iCAFs. The m8 population, exhibiting high levels of proliferation markers (Fig. 2E; *Mki67*, *Top2a*) and cell cycle-related signatures (Fig. 2F; E2F_targets, MYC_targets_V1), likely corresponds to a cycling CAF subpopulation (Fig. 2E). In contrast, we did not detect expression of any of apCAF markers in the CAF subpopulations (fig. S2B). We identified genes preferentially expressed in each CAF subpopulation (fig. S2C).

Examination of cell type-specific markers indicated that all populations of EpCAM^-^/CD45^+^ cells expressed *Adgre1* (which encodes F4/80) and *Cd68*, markers for macrophages (Fig. 2B). The m2 subpopulation expressed markers of M1 macrophages, *Il1b* and *Itgax* (*Cd11c*) (*30*), and showed elevated inflammatory signals (*30*) (fig. S2, D and E). The m5 subpopulation demonstrated characteristics of M2 macrophages, marked by expression of *Cd163*, *Mrc1* (*30*) and genes associated with oxidative phosphorylation (*31*) (fig. S2, D and E). The m7 subpopulation appeared to be a cycling macrophage population, as indicated by high levels of proliferation markers (fig. S2D; *Mki67*, *Top2a*) and cell cycle-related signatures (fig. S2E; E2F_targets, Mitotic spindle, G2M_checkpoint). We did not detect cell populations corresponding to other immune cell types, presumably due to their rarity and the compromised immunological environment in NOG mice.

Subsequently, we investigated whether CAFs or macrophages were localized near to KRT17-positive cancer cells. Co-immunostaining with Vimentin or F4/80 indicated that Vimentin^+^ cells, but not F4/80^+^ cells, were localized close to KRT17^+^ cells (Fig. 2G and fig. S2F). Close localization to KRT17^+^ cells were also observed for cells expressing Pdgfr, another CAF marker (fig. S2G). These data suggest the potential interaction between CAFs and the KRT17^+^ slow-cycling CSCs.

### KRT17^+^ cancer cells co-localize with myCAFs

To systematically examine the microenvironment surrounding the KRT17^+^ cancer cells, we assessed the localization of the identified tumor and non-tumor populations by integrating RNA-seq data with Visium spatial transcriptomics (*32, 33*). Transcriptomes from tissue sections of two CRC-6-derived xenograft tumors (CRC-6_T1, CRC-6_T2) were subjected to Visium analyses (CRC-6_T1, 1747 evaluated spots, 9820 median human UMIs/spot, 287 median mouse UMIs/spot; CRC-6_T2,1525 evaluated spots, 14318 median human UMIs/spot, 950 median mouse UMIs/spot). Based on gene expression profiles, the Visium spots were classified into ten clusters (c0–c9) in CRC-6_T1, and eight clusters (c0–c7) in CRC-6_T2 (Fig. 3A and fig. S3A).

**Fig. 3.**
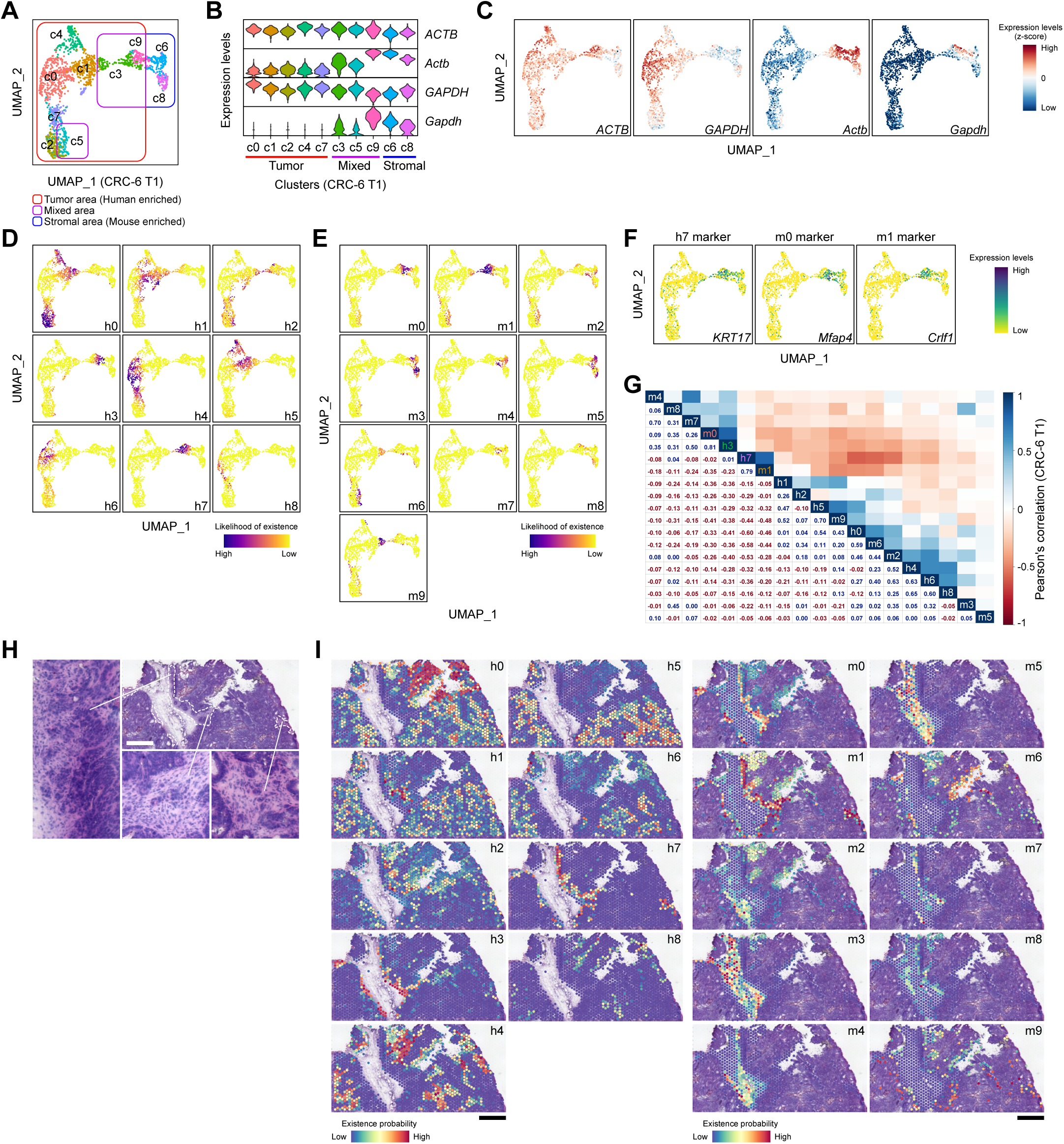
KRT17^+^ cancer cells co-localize with myCAFs. (**A**) UMAP plot of Visium spots of CRC-6_T1. Ten clusters are denoted by distinctive colors. Three types of clusters (a tumor-rich cluster, a stroma-rich cluster, and a mixed cluster) are marked by the indicated boxes. (**B**) Violin plots showing expression of human (*ACTB*, *GAPDH*) and mouse (*Actb*, *Gapdh*) housekeeping genes in the Visium clusters. (**C**) UMAP Feature plots of Visium spots showing expression of the housekeeping genes (CRC-6_T1). (**D, E**) Heatmap presentation of the prediction scores for the cancer cell subpopulations (h0–h8) (D) and the stromal cell subpopulations (m0–m9) (E) (CRC-6_T1). (**F**) UMAP Feature plots of Visium spots showing expression of specific markers for the h7 (*KRT17*), m0 (*Mfap4*), and m1 (*Crlf1*) subpopulation. (**G**) Correlation matrix showing the likelihood of the co-presence of the indicated cancer/stromal subpopulations in Visium spots (CRC-6 T1). Correlation coefficients between the indicated subpopulations are shown as calculated values and color gradients. (**H**) Hematoxylin-stained images of xenograft tumor (CRC-6 T2). Areas surrounded by dashed lines indicate stromal spaces, which are shown in detail in the magnified panels. Scale bar, 1 mm. (**I**) Heatmap presentation of the spatial distribution of the indicated subpopulations in xenograft tumor (CRC-6_T2). Scale bar, 1 mm.

Next, we assessed the spatial distribution of human tumor and mouse stromal cells in each cluster by using human (*ACTB*, *GAPDH*) and mouse (*Actb*, *Gapdh*) housekeeping genes as surrogate markers. We classified the clusters into three distinct groups based upon expression of the surrogate genes; tumor cell-rich clusters predominantly consisting of human tumor cells (CRC-6_T1: c0, c1, c2, c4, c7; CRC6_T2: c2, c3, c4, c5), stroma-rich clusters predominantly consisting of mouse non-tumor cells (CRC-6_T1: c6, c8; CRC6_T2: c0), and mixed clusters in which both human and mouse cells coexist (CRC-6_T1: c3, c5, c9; CRC6_T2: c1, c6, c7) (Fig. 3, B and C and fig. S3, B and C).

To evaluate the spatial localization of the tumor and non-tumor subpopulations, we calculated prediction scores for each subpopulation in each Visium spot and visualized them upon UMAP plots (Fig. 3, D and E and fig. S3D). Notably, the prediction scores for the *KRT17*^+^ cancer subpopulation (h7) indicated that this subpopulation predominantly localized in the mixed clusters (c3 and c9 in CRC-6_T1 and c6 and c7 in CRC-6_T2) (table S5 and S6). Consistent with this, cells expressing high levels of *KRT17* were observed in the same clusters in both tumors (Fig. 3F, fig. S3E, and table S7).

To evaluate co-localization of the tumor and non-tumor subpopulations on the same spot, we calculated co-localization coefficient indices based on the prediction scores for each subpopulation (CRC-6). Of note, the h7 subpopulation preferentially co-localized with the m1 subpopulation in CRC-6_T1 (Fig. 3G) and the m0 and m1 subpopulation in CRC-6_T2 (fig. S3F), indicating that KRT17^+^ slow-cycling cancer cells co-localize with myCAFs. Co-localization of the h7 subpopulation with the myCAF subpopulations (m0, m1) near the peripheral areas of tumors was visualized in a heatmap presentation of the prediction scores upon the tumor tissues (Fig. 3H and I, and fig. S3, G and H).

### CAF-derived TGF-β1 induces KRT17 expression in cancer cells

Co-localization of the KRT17^+^ subpopulation of cancer cells with myCAFs at the tumor periphery prompted us to explore the possibility that myCAFs might affect the metastatic capability of this subpopulation via diffusible signals. To identify the potential signaling pathways between these cells, we selected genes coding for diffusible ligands expressed by the myCAF subpopulations (m0 and m1). Ligand-receptor analyses (CellChat) (*34*) allowed us to select myCAF-expressed ligands whose receptor genes were highly expressed in the h7/h3 cancer subpopulations (Angptl4, Gas6, Spp1, TGF-β1-3, Tnfsf12, and Wnt5a) (Fig. 4A).

**Fig. 4.**
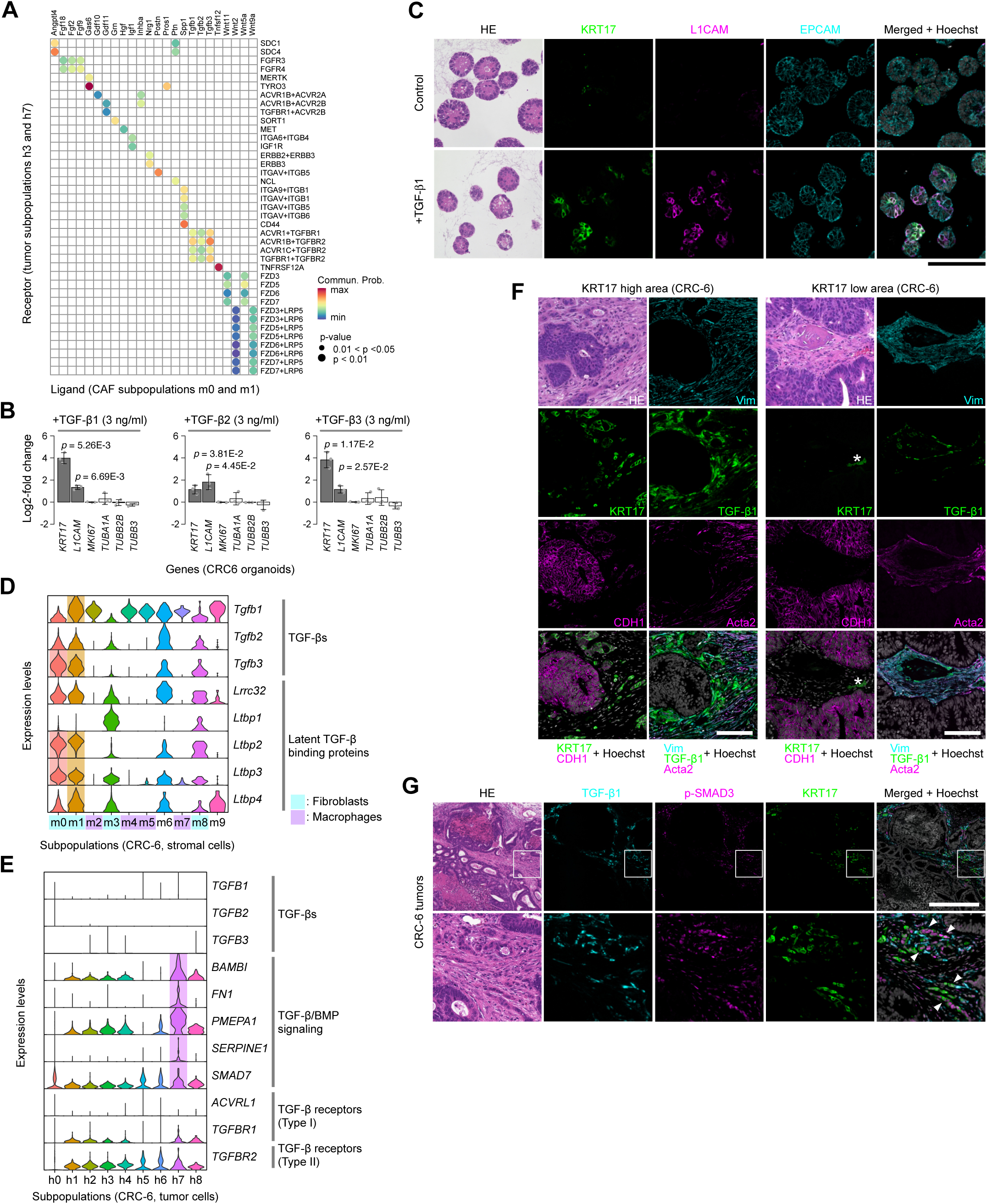
CAF-derived TGF-β1 induces KRT17 expression in cancer cells. (**A**) CellChat analysis of ligand-receptor interactions between myCAF-derived ligands and KRT17^+^ cancer cells-derived receptors. (**B**) Induction of h7 subpopulation-associated genes by TGF-β. Colon cancer organoids (CRC-6) were treated with the indicated isoforms of TGF-β1 (3 ng/mL) for 24 h, and induction of the indicated h7-associated genes and *MKI67* was evaluated by qPCR (n = 3). Genes induced with statistical significance (p < 0.05) are shown as gray bars. (**C**) Induction of *KRT17* and *L1CAM* in cancer organoids (CRC-6) treated with TGF-β. Organoids were treated with vehicle control or TGF-β1 (3 ng/mL) for 24 h, and used for immunostaining with the indicated antibodies or for H&E staining. Scale bar, 100 μm. (**D**) Violin plots showing expression of TGF-β isoform genes and latent TGF-β-binding proteins in stromal subpopulations. (**E**) Violin plots showing expression of genes involved in the TGF-β signaling pathway in cancer subpopulations (CRC-6). (**F**) Co-localization of KRT17^+^ cancer cells with TGF-β1^+^ CAFs in colon tumors. Xenograft tumor areas (CRC-6) populated with high or low numbers of KRT17^+^ cells were selected for immunostaining and H&E staining. Scale bar, 100 μm. (**G**) SMAD3 phosphorylation in KRT17^+^ cancer cells. The tumor/stromal border areas of xenograft tumors (CRC-6) were selected for immunostaining and H&E staining. Magnified images of the white boxes are shown at the bottom. Arrowheads indicate tumor cells expressing both KRT17 (cytoplasm) and phosphorylated SMAD3 (nuclei). Scale bar, 100 μm.

Among these selected ligands, TGF-β isoforms are of particular interest due to their known role in activating CAFs (*5, 9, 10*) and in enhancing metastatic capability (*6*). When we treated patient-derived organoids with TGF-β isoforms, we found that they upregulated some h7 subpopulation-specific genes, *KRT17* and *L1CAM* (Fig. 4B and fig. S4A), while none of the other ligands induced the h7 subpopulation-specific genes by more than 2-fold; the exception was Wnt5a, which slightly induced *KRT17* (fig. S4B). Immunostaining confirmed that the TGF-β1 treatment induced KRT17 and L1CAM in largely overlapping organoid cell populations (Fig. 4C and fig. S4C).

As expected from the receptor-ligand analyses, *Tgfb1-3* and genes encoding latent TGF-β-binding proteins were expressed preferentially in the m1 subpopulation of non-tumor cells (Fig. 4D). While expression of TGF-β receptors (*TGFBR1* or *TGFBR2*) was observed in most cancer subpopulations (Fig. 4E and fig. S4D), downstream targets of TGF-β signaling including *BAMBI* (*35*), *PMEPA1* (*36*) were induced preferentially in the h7 subpopulation (Fig. 4E and fig. S4D), indicating specific activation of the TGF-β pathway in this subpopulation.

Next, we performed immunostaining to examine whether myCAFs expressing TGF-β are located near KRT17^+^ cancer cells in colon tumors. In accordance with the Visium data, we observed that Vimentin^+^ CAFs expressing TGF-β1 co-localized with the KRT17^+^ cancer cells, but not with KRT17^-^ cancer cells (Fig. 4F and fig. S4E). We observed largely reciprocal staining of TGF-β1 and Acta2 (α-SMA) in Vimentin-positive CAFs, with Acta2^+^ cells located in areas relatively distant from KRT17^+^ cancer cells (Fig. 4F and fig. S4E). We did not observe significant TGF-β1 co-staining of F4/80^+^ macrophages (fig. S4F). In accordance, data mining of public single-cell datasets from clinical colon cancers (*14*) suggests that, while TGF-β is predominantly expressed in myeloid cells in nontumor tissues, it is preferentially expressed in stroma cells in tumor tissues (fig. S4G).

Immunostaining of tumor-stroma border areas revealed that a significant fraction of the KRT17^+^ cells that localized near to TGF-β1^+^ stromal cells expressed phosphorylated SMAD3, a main downstream regulator of TGF-β (Fig. 4G). In accordance, data mining of public ChIP-seq databases revealed that SMAD2/3 bind to the promoter regions of KRT17 in cancer cells (fig. S4H), suggesting that TGF-β1-dependent KRT17 induction is mediated via activation of SMAD2/3.

### Co-culture with CAFs generates KRT17^+^ migrating cancer cells

To recapitulate CAF-dependent generation of the KRT17^+^ cancer cells *in vitro*, we established primary fibroblasts from the xenograft tumors and examined whether co-culturing these mouse fibroblasts with cancer organoids induces KRT17 expression in cancer cells. Co-culturing CRC-6 or CRC-32-derived organoids with the mouse fibroblasts led to formation of cell aggregates with distinctive structures, in which fibroblasts were positioned centrally within the surrounding cancer organoids (Fig. 5, A and B) (*37*). Remarkably, immunostaining of the cancer-CAF aggregates revealed the presence of KRT17^+^ and L1CAM^+^ cancer cells that disseminated into the central fibroblast areas (Fig. 5B), whereas we did not observe the cell expressing KRT17 and/or L1CAM when the organoids were cultivated alone (Fig. 5B). Accordingly, co-culture with CAFs induced expression of the h7 subpopulation-associated genes, including *KRT17* and *L1CAM* (Fig. 5C). By contrast, we did not observe detectable expression of these genes after co-culture with xenograft tumor-derived macrophages (fig. S5A).

**Fig. 5.**
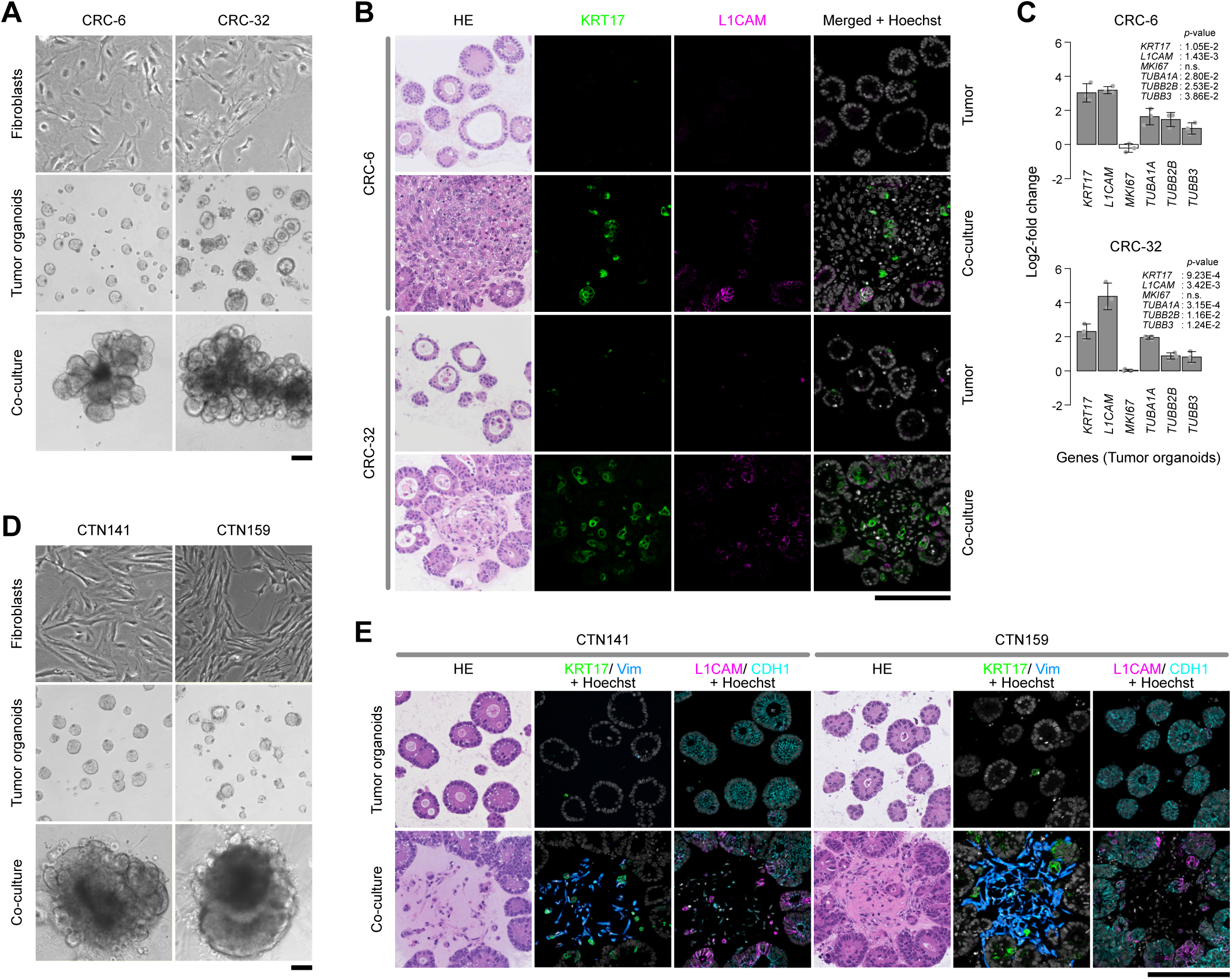
Co-culture with CAFs generates KRT17^+^ migrating cancer cells. (**A**) Establishment of *in vitro* co-cultures of human cancer organoids and mouse CAFs (CRC-6 and CRC-32). Phase contrast images of cancer organoids and matched xenograft-derived mouse CAFs that were cultivated alone or co-cultivated for 7 days. Scale bar: 100 μm. (**B**) Cancer organoids grown in the presence or absence of mouse CAFs for 7 days were used for H&E staining (left panels) or for immunostaining with the indicated antibodies. Scale bar: 100 μm. (**C**) CAF-dependent induction of h7-associated genes in cancer organoids. Cancer organoids shown in B were used for qPCR to detect the indicated genes; qPCR values obtained in the absence or presence of matched CAFs were compared. Genes induced with statistical significance (p < 0.05) are shown as gray bars. n = 3. (**D**) Colon cancer organoids and CAFs that were simultaneously established from surgical specimens (CTN141, CTN159) were cultivated alone or co-cultivated for 7 days. Phase contrast images of cells are shown. Scale bar: 100 μm. (**E**) The cancer organoids grown in the presence or absence of matched human CAFs for 7 days were used for H&E staining (left panels) or immunostaining with indicated antibodies. Scale bar: 100 μm.

To determine whether CAFs derived from human cancer also induce expression of h7 subpopulation-associated genes, we simultaneously established patient-derived cancer organoids and CAFs from clinical specimens in which both KRT17 and L1CAM were expressed (Fig. 1J, fig. S1K, and table S3) and used them in the co-culture experiments. Co-culture with the matched human CAFs led to formation of aggregates with a concentric structure (Fig. 5D and fig. S5B). Again, we observed emergence of KRT17^+^ and/or L1CAM^+^ disseminating cancer cells in the presence of CAFs (Fig. 5E and fig. S5C). These data suggest that CAFs induce the KRT17^+^/L1CAM^+^ cancer cells with invasive capability *in vitro*.

### KRT17 is essential for development of invasive cancer cells with metastatic capability

To examine the functional role of KRT17 during CAF-dependent invasion, we evaluated dissemination of cancer cells within the co-cultured aggregates (Fig. 5 and fig. 5) as a proxy for cancer invasion. We conducted CRISPR knockout of *KRT17* in cancer cells (Fig. 6A and fig. S6A) and measured the KRT17-deficient cells in the co-culture assays. *KRT17* knockout effectively eliminated invasion of cancer cells into the fibroblast-dominated areas (Fig. 6, B and C, and fig. S6, B and C).

**Fig. 6.**
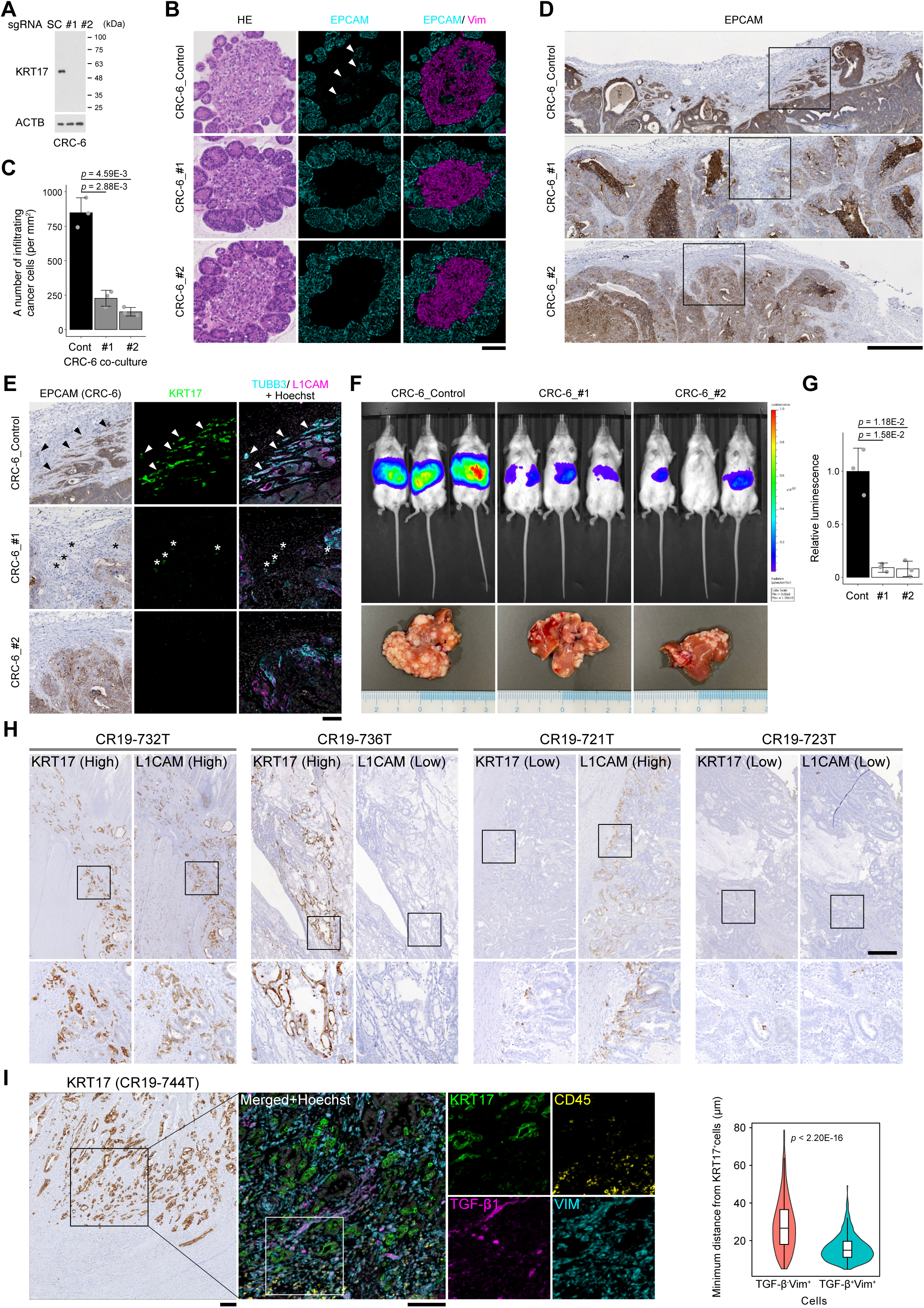
KRT17 is essential for development of invasive cancer cells with metastatic capability. (**A**) CRISPR/Cas9-mediated knockout of *KRT17* in cancer cells. Cancer spheroids (CRC-6) expressing Luciferase (CRC-6/luc2-BFP) were introduced with sgRNAs targeting KRT17 (sgKRT17#1, sgKRT17#2) or non-targeting scramble control sgRNA (Control) and subjected to single-cell FACS sorting and expansion. Cancer spheroids introduced with the indicated sgRNAs were treated with 3 ng/mL TGF-β1 for 24 h and used for western blot with the indicated antibodies. (**B**) *In vitro* co-culture with KRT17-deficient cancer cells. Cancer spheroids described in A were converted into organoids and used for coculture assays with mouse CAFs. Formed aggregates were subjected to H&E staining and to immunostaining with the indicated antibodies. EpCAM-positive cancer cells infiltrating into Vimentin-positive stromal areas are indicated by arrowheads. Scale bar: 100 μm. (**C**) The numbers of the infiltrating cancer cells was calculated (n = 3). (**D**) Reduced stromal infiltration of the KRT17-deficient cancer cells. Xenograft tumors generated from the control and KRT17-deficient spheroids were subjected to immunostaining with an anti-EpCAM antibody. Scale bar: 500 μm. (**E**) The boxed areas shown in D were immunostained with the indicated antibodies. Arrowheads indicate invasive tumor cells. Asterisks: autofluorescence of red blood cells. Scale bar: 100 μm. (**F**) (upper panels) Bioluminescence images of luciferase activity in metastasized livers, measured at 6 weeks after splenic injection of control or KRT17-deficient cells into NOG mice; (lower panels) representative images of metastasized livers. (**G**) Normalized luminescence values calculated by measuring total flux in the metastasized livers shown in F (n = 3). (**H**) Immunostaining of human colon cancer specimens. Thirty cases of FFPE samples (Stages I–III) were subjected to IHC staining with anti-KRT17 or anti-L1CAM antibodies. Representative staining images from four groups (KRT17^high^/L1CAM^high^, KRT17^high^/L1CAM^low^, KRT17^low^/L1CAM^high^, KRT17^low^/L1CAM^low^) are shown. Magnified images of the black boxes are shown in the bottom panels. Scale bar: 500 μm. (**I**) Co-localization of TGF-β1^+^ CAFs with KRT17^+^ cancer cells in human cancers. KRT17^+^ specimens, shown in the left column of the immunostained images (black box) were subjected to immunofluorescence analyses with the indicated antibodies. Magnified images of the while boxes were shown in the right-hand panels. Scale bars: 100 μm. Violin plots of the minimum Euclidean distances from KRT17⁺ cells to TGF-β^-^Vim⁺ and TGF-β^+^Vim⁺ cells are shown in the rightmost panel.

Next, we utilized the *KRT17* knockout cells to assess whether KRT17 is required for invasion and metastasis of xenograft tumors. While the KRT17 knockout did not significantly affect tumor growth (fig. S6, D and E), it led to a significant reduction in the number of EpCAM^+^ invasive cells (Fig. 6D and fig. S6F), many of which expressed L1CAM and/or TUBB3 (Fig. 6E and fig. S6, G and H). Moreover, the knockout substantially reduced liver metastasis following splenic injection of the cancer cells (Fig. 6, F and G).

### KRT17^+^ cancer cells co-localize with TGF-β1^+^ CAFs in clinical colon cancers

Next, we examined whether expression of KRT17 and L1CAM is associated with advanced stages of human colon cancer. Immunostaining of surgical FFPE specimens from 30 cases (table S8) classified colon cancer into four groups based upon expression of KRT17 and L1CAM (Fig. 6H, KRT17^high^/L1CAM^high^ (9 cases), KRT17^high^/L1CAM^low^ (6 cases), KRT17^low^/L1CAM^high^ (6 cases), KRT17^low^/L1CAM^low^ (9 cases)). Indeed, tumors expressing high levels of both KRT17 and L1CAM were associated with advanced stages (fig. S7A).

Subsequently, we examined the clinical relevance of KRT17 induction by CAF-derived TGF-β by performing immunofluorescence analyses of these FFPE specimens. We preferentially observed TGF-β1^+^/Vimentin^+^ cells in KRT17^high^/L1CAM^high^ and KRT17^high^/L1CAM^low^ cases, but not in KRT17^low^/L1CAM^high^ and KRT17^low^/L1CAM^low^ cases (Fig. 6I and fig. S7B). In the KRT17^high^/L1CAM^high^ cases, TGF-β1^+^/Vimentin^+^ cells were found in tumor regions but not in neighboring non-tumor regions (fig. S7B, right panels). Thus, TGF-β-expressing CAFs in clinical tumors colocalize specifically with KRT17^+^ colon cancer cells.

### The KRT17^+^ cells exhibit hallmarks of senescence and are eliminated by senolytic treatment

We observed that some KRT17^high^ cells exhibited enlarged, irregularly shaped cells bodies, reminiscent of morphological features of senescent cells (Fig. 7A) (*38*). Gene expression profiles of KRT17^high^ cells displayed multiple hallmarks of cell senescence (*38, 39*), including cell cycle inhibition as indicated by reduced *MKI67* expression (Fig. 1B and fig. S1B) and elevated expression of key senescence associated secretory phenotype (SASP) components, including *CXCL1*, *CXCL8*, and *TNF* (Fig. 1G and fig. S1F). In addition, gene set enrichment analyses (GSEA) indicated that the h7 cancer subpopulation exhibits a senescence-associated transcriptional profile (Fig. 7B). Moreover, this subpopulation showed elevated expression of *CDKN2A*, a senescence-associated cyclin-dependent kinase inhibitor encoding p16^INK4a^, and *BCL2L1*, a member of BCL2 family that promotes survival of senescent cells (Fig.7C). In accordance, we observed extensive co-localization of KRT17 with BCL2L1 and CDKN2A at invasion fronts (Fig. 7D).

**Fig. 7.**
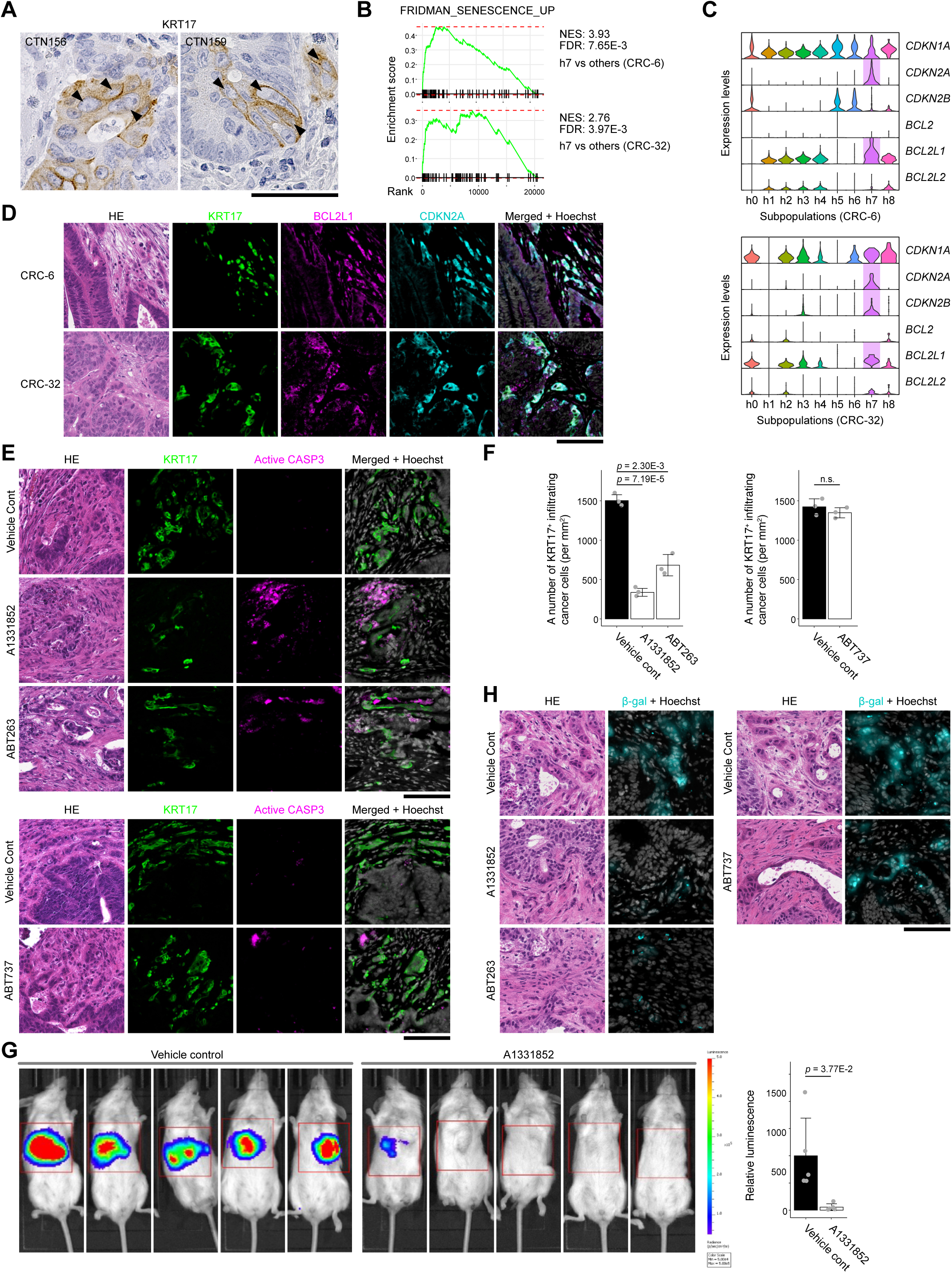
The KRT17^+^ cells exhibit hallmarks of senescence and are eliminated by senolytic treatment. (**A**) Magnified images of immunostained KRT17^+^ cells located at the invasion front of human colon cancer (CTN156 and CTN159). Arrowheads show irregularly shaped KRT17^+^ cells. Scale bar: 50 μm. (**B**) GSEA evaluation of a senescence signature (FRIDMAN_SENESCNCE_UP) in the h7 cancer subpopulation of xenografted tumors in comparison to the other subpopulations (CRC-6 and CRC-32). (**C**) Violin plot showing gene expression of cyclin-dependent kinase inhibitors and BCL2 family members in cancer subpopulations. (**D**) H&E staining and immunostaining of the xenograft tumors (CRC-6 and CRC-32) with the indicated antibodies. Scale bar: 100 μm. (**E**) H&E staining and immunostaining of the xenograft tumors (CRC-6) following senolytic treatments. Scale bar: 100 μm. (**F**) Quantitative evaluation of the stroma-infiltrating KRT17^+^ cells shown in E. The number of infiltrating cells were calculated as (the number of KRT17^+^ cells in the stromal area populated within Vimentin^+^ cells) / (area size). Average data for five representative area are shown (*n* = 3). (**G**) Bioluminescence images of luciferase activity in metastasized livers. NOG mice subjected to splenic injection of CRC-6 cells were orally administered with A1331852 or vehicle control for two weeks before the bioluminescence assay. Normalized luminescence values are shown in the right panel (n = 5). (**H**) Fluorogenic staining of SA-beta-gal-positive cells in the xenografted tumors shown in D. Scale bar: 100 μm.

Senescent cells exert detrimental effects in variety of physiological and pathological contexts, and their targeted elimination using senolytic agents has garnered significant interest for therapeutic applications (*40, 41*). Among the key target of senolytics are BCL2 family proteins, which functions as anti-apoptotic regulators. Given the elevated expression of BCL2L1 in our model, we aimed to eliminate the KRT17^+^ senescent cells using senolytic agents.

To assess the efficacy of BCL2L1-targeting senolytics on the KRT17^+^ senescent cells, we administered these agents in xenografted mice. Treatment with A1331852 (*42*), ABT263 (*43*), or ABT737(*44*), senolytic agents, did not significantly affect growth of xenografted tumors (fig. S7C). Remarkably, treatment with A1331852 (*42*) or ABT263 (*43*) resulted in significantly reduction in KRT17^+^ cells at the invasion fronts, whereas treatment with ABT737, which mainly targets BCL2, had minimal effect (Fig. 7, E and F). Consistently, we observed a marked increase of apoptotic cells expressing active CASP3 following treatment with A1331852 or ABT263 (Fig.7E), indicating that senolytic therapy induced apoptotic cell death in KRT17^+^ senescent cells. Furthermore, treatment with A1331853 markedly reduced the extent of metastasis after splenic injection of cancer cells (Fig. 7G), indicating that the senolytic treatment also blocks post-entry process of liver metastasis.

Next, we examined the presence of cells expressing active senescence-associated β-galactosidase (SA-β-gal), a widely used marker of cell senescence (*38, 45*), in the xenografts. As expected, β-gal^+^ cells were observed at the invasion front of untreated xenografts. Notably, treatment with A1331852 or ABT263, but not ABT737, largely eliminated β-gal^+^ cells (Fig. 7H). As expected, KRT17 knockout also caused a marked reduction of the β-gal^+^ cells (fig. S7D). Collectively, these findings demonstrate that the senolytic treatment effectively eliminates KRT17^+^ invasive cancer cells with senescent phenotypes.

Finally, data mining of public single-cell datasets used in Fig. S4G revealed that colon cancer epithelial cells harbor a subpopulation expressing elevated levels of *KRT17*, *BCL2L1*, and *CDKN2A* (c2 subpopulation, fig. S7E), suggesting a widespread presence of KRT17^+^ colon cancer cells with a senescent phenotype.

## DISCUSSION

In this study, we combined single-cell analyses, spatial transcriptomics, and functional assays using patient-derived colon cancer models to investigate how tumor-stroma interactions contribute to the emergence of invasive cancer cell states. Our data support a model in which a KRT17^+^ slow-cycling, CSC-like subpopulation is induced at tumor-stroma interfaces through CAF-derived TGF-β signaling. This spatially restricted induction provides a framework for understanding how microenvironmental cues may promote the emergence of metastasis-initiating cell-like states. The functional importance of KRT17^+^ cells in invasion and metastasis observed in our experimental models is consistent with previous reports linking high KRT17 expression to poor prognosis in colorectal cancers (*46*) and several other cancers (*47*).

Notably, the KRT17^+^ cells exhibit features reminiscent of cellular senescence. From a therapeutic perspective, these findings had important implications: Senolytic therapies, designed to selectively eliminate senescent cells, have gained interest as potential cancer treatments (*40, 41*). Since a variety of anti-cancer therapies induce senescence, multiple clinical trials are evaluating senolytic agents in combination with conventional treatments (*40*). From a therapeutic perspective, our findings raise the possibility that senolytic strategies targeting anti-apoptotic BCL2 family proteins may preferentially eliminate KRT17^+^ cells and thereby suppress metastatic progression.

Cellular senescence plays complex roles in tumorigenesis, exerting both tumor-suppressive and tumor-promoting effects (*45, 48*). While initially acting as a barrier to tumor progression, senescence can paradoxically enhance cancer progression through non-cell-autonomous mechanisms, primarily via the senescence-associated secretory phonotype (SASP). SASP factors have been implicated in priming cancer cells for re-proliferation and relapse by conferring stemness-like properties (*48, 49*). Our findings that show the KRT17^+^ invasive cells exhibit phenotypic resemblance to senescence cells broaden the recognized impact of senescence in cancer progression and provide another rationale for targeting senescent cancer cells in cancer therapy.

Another key finding is the identification of a specific CAF subset (m1 myCAFs) as a major source of TGF-β in our model. While TGF-β has long been implicated in metastasis, our data provide a spatial and cellular context suggesting that myCAF-derived TGF-β may contribute to the induction of a senescence-associated invasive state in adjacent cancer cells. This highlights functional heterogeneity among CAF populations: whereas iCAFs have been linked to therapy resistance (*13*), our results demonstrate that myCAFs can actively promote cancer progression. Targeting CAF subpopulations or modulating their signaling outputs may therefore represent an additional strategy to disrupt MIC generation. Overall, our study highlights the value of integrating single-cell and spatial transcriptomic approaches with functional validation to uncover context-dependent cancer cell states (*32, 33*). Applying similar strategies across tumor types may help identify conserved or context-specific mechanisms underlying metastasis and therapeutic resistance.

### Limitation of our study

Our study has several limitations. First, functional analyses *in vivo* were performed using xenograft models in immunodeficient mice. These systems do not fully recapitulate the complexity of the human tumor microenvironment, particularly the role of adaptive immunity. Therefore, validation in immunocompetent or humanized models will be necessary to confirm the generalizability of CAF-mediated KRT17 induction.

Second, although we demonstrate that KRT17 is functionally required for invasive behavior and metastasis, the downstream molecular pathways through which KRT17 orchestrates the metastasis-initiating cell state remain to be fully elucidated. Whether KRT17 acts primarily as a structural scaffold or participates in signaling modulation is an important question for future mechanistical studies.

Third, while KRT17 cells exhibit core hallmark of cellular senescence, including expression of CDKN2A and BCL2L1 and senescence-associated gene signatures, these cells may represent a plastic, senescent-like transition state. The physiological depth of this senescence state requires further characterization.

Finally, while senolytic agents effectively eliminated KRT17 cells and suppressed metastasis in our models, their clinical applicability requires caution. Future studies should evaluate the safety profiles of these strategies in clinically relevant settings and investigate potential effects on normal tissues and immune responses.

## ACKNOWLEDGMENTS.

Human cancer specimens were provided by the National Cancer Center Biobank. We thank Yoshie Okimoto for technical assistance, National Cancer Center Research Institute Core Facility for preparation of the samples used for Immunohistochemistry, and the Division of Medical Research Support, Advanced Research Support Center (ADRES), Ehime University, for their technical assistance. This research was supported by Grant-in-Aids for Scientific Research from the Japan Society for the Promotion of Science (JSPS) under grant numbers JP21H03081 (K.O.) and 18K07283 (D.S.); the Japan Agency for Medical Research and Development (AMED) under Grant numbers JP19cm0106563h0001 and JP21cm0106183h0001 (K.O.); the Japan Science and Technology Agency (JST) for CREST (Core Research for Evolutional Science and Technology) under grant number 21-211035300 (K.O.); the Japan Health Research Promotion Bureau (JH) under grant number 2020-B-03 (K.O.); The Vehicle Racing Commemorative Foundation (K.O. and M.M.); Princess Takamatsu Cancer Research Fund (D.S.); and Single Cell Medical Network (RIKEN IMS) (K.O.).

## MATERIALS AND METHODS

### Clinical samples

Clinical materials that were derived from surgical specimens of colon cancer and obtained with written informed consent were obtained from the Biobank of National Cancer Center. Surgical specimens were subjected to FFPE analyses or used to establish patient-derived cells. Clinicopathological information is presented in table S3 and S8. All procedures were performed using protocols reviewed and approved by the Ethics Committee of the National Cancer Center (#2008-097), Teikyo University (#22-020-3), and Ehime University Hospital (#2023A063).

### Antibodies

Antibodies for ACTB (# 3700), KRT17 (#12509), L1CAM (#90269), Vimentin (#5741), F4/80 (#70076), EPCAM (#14452), CDH1 (# 3195), CD45 (#13917), BCLXL (#2764), and cleaved Caspase 3 Asp175 (# 9661) were purchased from Cell Signaling Technology. Antibodies for TGF-β1 (ab215715) and ACTA2 (ab7817) were purchased from Abcam. An antibody for CDKN2A was purchased from Santa Cruz Biotechnology. Anti-human EPCAM APC-conjugated was purchased from BD Biosciences. Antibodies for Cd16/32 (#101302) and TUBB3 (# 801202) were purchased from BioLegend. An antibody for MKI67 (#M7240) was purchased from Agilent. Anti-mouse Cd45 PE-conjugated was purchased from TONBO Biosciences. An antibody for PROX1 (AF2727) was purchased from R&D Systems. Anti-mouse Cd45 PE-conjugated was purchased from TONBO Biosciences. Secondary antibodies, donkey anti-Goat IgG, Alexa Fluor 555-conjugated (A21432), donkey anti-mouse IgG, Alexa Fluor 488-conjugated (A21202), donkey anti-mouse IgG, Alexa Fluor 555-conjugated (A31570), donkey anti-mouse IgG, Alexa Fluor 647-conjugated (A31571), donkey anti-rabbit IgG, Alexa Fluor 555-conjugated (A31572), donkey anti-rabbit IgG, and Alexa Fluor 647-conjugated (A31573) were purchased from Invitrogen. A secondary antibody, donkey anti-rabbit IgG, Alexa Fluor 750-conjugated (ab175728), was purchased from Abcam.

### Chemicals and Reagents

The catalog numbers and vendors for chemicals and reagents are shown as the following: Collagenase P (Roche, #11249002001), Dispase II (Roche, #492078001), DNase I (Roche, #11284932001), Calcein AM (Invitrogen, C3100MP), StemPro hESC SFM (Gibco, A1000701), MEM-α (Fuji Film, #135-15175), DMEM/F12-GlutaMAX (Gibco, #10565018), HEPES 1M (Gibco, #15630080), N-2 Supplement (Gibco, #17502-048), B-27 supplement (Gibco, #17502-048), N-acetyl-L-cysteine (Sigma-Aldrich, A7250), 30% (w/v) BSA solution (Fuji Film, #017-22231), Insulin, Transferrin, Selenium Solution (Gibco, #41400045), HBSS (Gibco, #14025092), D-PBS(-) (Fuji Film, #045-29795), Accumax (Innovative cell technology, AM105), TripLE Express (Gibco, #12604013), Matrigel basement membrane matrix (Corning, #354234), Matrigel growth factor reduced (Corning, #356231), Cell Recovery Solution (Corning, #354253), Tyramide Super Boost kit Alexa Fluor 488 (Thermo Fisher Scientific, B40922), Alexa Fluor 488 Tyramide reagent (Thermo Fisher Scientific, B40953), Alexa Fluor 555 Tyramide reagent (Thermo Fisher Scientific, B40955), Alexa Fluor 647 Tyramide reagent (Thermo Fisher Scientific, B40958), A1331852 (Selleck, S7801), ABT263 (Selleck, S1001), and ABT737 (Selleck, S1002).

### Recombinant proteins

The catalog numbers and vendors for recombinant proteins are shown as the following: Human recombinant EGF (Gibco, PHG0311), Human recombinant FGF-2 (Gibco, PHG0021), Mouse recombinant Tgf-β1 (BioLegend, #763104), Mouse recombinant Tgf-β2 (PeproTech, #100-35B), Mouse recombinant Tgf-β3 (PeproTech, #100-36E), Mouse recombinant Angptl4 (Sino Biological, #50356-M07H), Mouse recombinant Gas6 (Sino Biological, #58026-M08H), Mouse recombinant Spp1 (BioLegend, #763606), Mouse recombinant Tnfsf12 (Sino Biological, #50174-M01H), and Human/Mouse recombinant Wnt5a (R&D Systems, P22725).

### Culture of cancer spheroids

Establishment of cancer spheroids from surgical specimens was reported previously (*50*). Colon cancer spheroids were grown on ultralow attachment culture dishes (Corning, #3471) in Spheroid medium (StemPro hESC SFM (Gibco, A1000701) supplemented with 8 ng/mL FGF-2 (Gibco, #13256-029), 20 μM Y-27632 (Fuji Film, #036-24023), and penicillin-streptomycin (Gibco, #15140-122)).

### Establishment of patient-derived organoids

To generate colon cancer organoids from surgical specimens, tumor tissues (approximately 5 mm³) were washed with ice-cold D-PBS(−), minced with a razor blades, and incubated for 45 min at 37°C in 1 mL of Digestion buffer (HBSS (Gibco, #14025092) containing 3 μg/mL collagenase P (Roche, #11249002001), 1 μg/mL Dispase II (Roche, #4942078001), 1 μg/mL DNase I (Roche, #11284932001), and 1 μM dithiothreitol (Sigma-Aldrich, #43815-1G)). The dissociated tumor cells were passed through a 100 μm filter, washed twice with FACS buffer (D-PBS(−) containing 0.5% BSA and 0.5 mM EDTA), and then subjected to cell culture of cancer organoids as described in the following section.

### Cell culture of cancer organoids

The dissociated organoid cells were suspended in 500 μL of E medium (DMEM/F12-GlutaMAX (Gibco, #10565-018) supplemented with penicillin-streptomycin (Gibco, #15140-122), 10 mM HEPES (Gibco, #156-30080), Insulin-Transferrin-Selenium (Gibco, #41400-045), N-2 (Gibco, #17502-048), B-27 (Gibco, #12587-010), 1 mM N-acetylcysteine (Sigma-Aldrich, A7250), 50 ng/mL human EGF (Gibco, PHG0311)) supplemented with 10 μM Y27632 (Fuji Film, #036-24023), and then cultured overnight in 24-well plates coated with 50 μL of Matrigel (Corning, #354234). Subsequently, the attached cells were overlaid with 50 μL of Matrigel, followed by adding 500 μL of E medium supplemented with 10 μM Y27632. The medium was replaced every 2 days. For splitting cells, the organoids were digested with TrypLE Express (Gibco, #12604013) and re-plated once a week.

### Conversion of cancer spheroids into organoids

To convert colon cancer spheroids into organoids, the spheroids were dissociated with Accumax and incubated under organoid conditions. Next, 5×10^3^ of the dissociated cells were suspended in 500 μL of E medium supplemented with 10 μM Y27632 and seeded overnight onto 24-well plates coated with 50 μL of Matrigel (Corning, #354234). Subsequently, the attached cells were overlaid with 50 μL of Matrigel followed by adding 500 μL of E medium supplemented with 10 μM Y27632.

### Preparation of CAFs

To prepare CAFs from surgical samples or xenograft tumors, tumor tissues were enzymatically dissociated with the digestion buffer, as described in the Establishment of patient-derived organoids section. Subsequently, the dissociated cells were suspended in 10 mL of MEMα (Fuji Film, #135-15175) containing 10% FBS and penicillin-streptomycin and seeded onto 10-cm culture dishes. The culture medium was replaced every 2 days. The cells grown to achieve ∼80% confluency were then dissociated with 0.25% trypsin-EDTA (Gibco, #25200056) and replated at a 1:5 dilution.

### Preparation of macrophages from cancer tissues

Mouse xenograft tumors (CRC-6) were enzymatically dissociated as described in the Preparation of CAFs section. The dissociated cells were suspended in 10 mL of Macrophage medium (Advanced RPMI 1640 (Gibco, #12633012) containing 10% FBS and penicillin-streptomycin) supplemented with a x1/10 volume of conditioned medium from L cells (ATCC, CRL-2648) as previously described (*51*), and then seeded onto 10 cm non-coated plastic dishes (Eiken Chemical, AU3240). The conditioned medium was prepared according to the protocol provided by ATCC (https://www.atcc.org/products/crl-2648). The culture medium was replaced every 2 days. The cells grown to achieve ∼80% confluency were detached from the dish by incubating with D-PBS(-) at 37°C and replated at a 1:5 dilution.

### Generation of mouse xenograft tumors

All animal experiments were performed in accordance with protocols that were reviewed and approved by the animal care and use committee of the National Cancer Center. To generate xenograft tumors, 5×10^5^ of Accumax-treated spheroid cells (CRC-6 or CRC-32) were suspended in 50 μL of a 1:1 mixture of Matrigel (Corning, #354234) and D-PBS(−) (Fuji Film, #045-29795) and subcutaneously transplanted into 8–12-week-old female NOG mice. Xenograft tumors were harvested when the longest diameter of the tumors reached approximately 5–10 mm. Tumor volumes were calculated using a standard formula (length x width x height x π/6).

### Treatment of xenograft tumors with senolytic agents

All senolytic agents were dissolved in the Senolysis buffer (60%Phosal50PG, 30%PEG400, and 10%EtOH) (*52*). One week after subcutaneous transplantation of dissociated spheroid cells (CRC-6), NOG mice were administered senolytic agents or the senolysis buffer for two weeks. A1331852 (25mg/kg) or ABT263 (50mg/kg) was orally administered five days per week, while ABT737 (25mg/kg) was intraperitoneally injected twice a week. Subsequently, xengrafted tumors of the treated mice were subjected to immunostaining of FFPE samples or fluorogenic staining of SA-β-gal.

### Fluorogenic staining of Senescence-associated β-galactosidase (SA-β-gal)

Xenograft tumors obtained from senolytic agent-treated or control mice were embedded in OCT compound (Tissue-Tek, #4583), snap-frozen on dry ice, and stored at −80°C. Frozen tissues were cryosectioned at a thickness of 7 μm, mounted on slides, and fixed with 10% formalin for 10 minutes. After permeabilization with 0.1% Triton X-100 for 10 minutes, SA-β-Gal activity was detected using the SPiDER-βGal fluorogenic substrate (Dojindo, #SG02) according to the manufacturer’s instructions. Nuclear DNA was counterstained with 10 μg/mL Hoechst 33342 (Invitrogen, H3570). Fluorescent images were acquired using a BZ-X810 fluorescence microscope (Keyence). For the quantification of cancer cells infiltrating stromal spaces, the number of KRT17^+^ cells located within areas populated with Vimentin^+^ cells was counted using ImageJ2 v2.14.0 (https://imagej.net/software/fiji/) (*53*).

### Treatment of cancer organoids with ligands

Colon cancer spheroids were dissociated with Accumax, and the dissociated cells (5×10^3^ cells/well) were seeded for 3 days onto 24-well plates containing Growth Factor-Reduced Matrigel (Corning, #356231) under the organoid culture conditions. The organoids were then incubated in E medium supplemented with ligands for an additional 24 hours. Cells were collected using Cell recovery solution (Corning, #354253) and subjected to immunostaining analyses or qPCR analyses.

### Co-culture experiments of patient-derived cancer organoids with CAFs

Dissociated cancer cells (5×10^3^ cells/well) were grown in 24-well plates containing Growth Factor-Reduced Matrigel (Corning, # 356231) under organoid conditions, either alone or together with equal number of dissociated CAFs. The medium was replaced every 2 days.

At 7 days after the initiation of co-culture, cells were harvested and subjected to immunostaining analyses or qPCR analyses. To quantify infiltration of cancer cells into the CAF-dominated central area, the number of EpCAM-positive tumor cells located inside Vimentin-positive areas was counted by ImageJ2 v2.14.0.

### Quantitative PCR (qPCR) analyses

Gene expression profiles of colon tumor organoids treated with various CAF-derived ligands, or co-cultured with CAFs, were analyzed by quantitative PCR (qPCR) using Thunderbird Cyber qPCR Mix (Toyobo, QPS-201). Cells under organoid culture were collected using Cell Recovery Solution (Corning, #354253) and total RNA was isolated using the NucleoSpin RNA Kit (Takara, #740955-250). The purified RNA was used to generate cDNAs using the Superscript VILO cDNA Synthesis Kit (Invitrogen, #11754-250). Real-time PCR analyses were performed on the StepOnePlus Real-Time PCR System (Applied Biosystems). Human-specific primers were designed by using Primer-BLAST software (http://www.ncbi.nlm.nih.gov/tools/primer-blast/) and listed in table S9. Quantification of gene expression was performed based on the ΔΔCt method, using the Ct values of *EPCAM* as a reference.

### FACS sorting of colon cancer cells from xenograft tumors

Xenograft tumors were minced and incubated in digestion buffer as described above. The dissociated cells were washed twice with FACS buffer and incubated with Accumax reagent (Innovative Cell Technologies, AM-105) for an additional 15 min at 37°C. The enzymatically digested cells were washed twice with FACS buffer and passed through a 100 μm filter. After blocking Fc receptors with an anti-mouse CD16/32 antibody (1:300; BioLegend, #101302) for 10 min on ice, the cells were stained for 30 minutes on ice with anti-mouse CD45 antibody conjugated to PE (1:300; TONBO biosciences, #50-0451) and an anti-human EpCAM antibody conjugated to APC (1:300; BD, #347200). The cells were then stained with Calcein AM dye (Invitrogen, C3100MP) for 15 min at 37°C, collected by centrifugation, resuspended in FACS buffer containing DAPI (100 ng/ml, Sigma-Aldrich, D9542), and sorted using a FACS AriaIIIu cytometer (Becton Dickinson). Viable cells were selected as Calcein AM-positive/DAPI-negative cells, and tumor cells were sorted as EpCAM-positive/CD45-negative cells. EpCAM-negative stromal cells were further sorted to collect CD45-positive and CD45-negative cells.

### Single-cell RNA-seq

The FACS-sorted cells (5×10^3^) were loaded onto a Chromium Next GEM Chip G (10x Genomics, PN-2000177) and captured on a Gel Beads-in-emulsion (GEMs) using a Chromium Controller (10x Genomics). Next, cDNA synthesis and sample-indexed library construction were performed with a Chromium Next GEM Single Cell 3’ Reagent Kit (10x Genomics, PN-1000121). The cDNA libraries were subjected to next-generation sequencing on the HiSeq2500 platform (Illumina). Mapping of raw sequence reads to the human GRCh38 and mouse mm10 reference genomes (version 2020-A, provided by 10x Genomics) and initial quality control were performed by using Cell Ranger software v3.0.2 (10x Genomics). For single-cell analyses of human cancer cells or mouse stromal cells, cells for which >70% of the reads mapped to the human or mouse reference genome, respectively, were used for further analyses. The UMI count matrices were used to select cells that meet the following criteria (cells with <15% mitochondrial gene counts; cells with <8,000 unique feature counts; and cells with >1,500 unique feature counts) using Seurat software v5.0.3 (https://satijalab.org/seurat/) (*54*) running on R V4.3.1. (https://www.r-project.org/).

### Analysis of single-cell RNA-seq data

Analyses of the filtered gene count matrices were performed using the Seurat pipeline as described previously (*55*). Data obtained from two replicates of the xenograft tumors were merged and without batch effect corrections. Single-cell gene expression counts were log-normalized using NormalizeData() with a scaling factor of 10,000. The top 2,000 most variable genes were identified using FindVariableFeatures(), scaled with ScaleData(), and subjected to principal component analysis (PCA) using RunPCA(). The dimensionality of the datasets was determined based on the percentage of variance explained by the top-ranked principal components. Cells were stratified using FindNeighbors() and FindClusters() functions with the following parameters: for cancer cells (CRC-6), dims = 1:6, resolution = 0.40; for cancer cells (CRC-32), dims = 1:9, resolution = 0.47; for stromal cells (CRC-6), dims = 1:9, resolution = 0.30. The resulting single-cell data was subjected to non-linear dimensional reduction using the UMAP method and projected in UMAP spaces using the RunUMAP() function. Integration of CRC-6 and CRC-32 datasets was performed using the CCAintegration method to identify clusters conserved across the tumors. Gene expression levels were represented as violin plots using VlnPlot() or as feature plots using FeaturePlot(). Potential ligand-receptor interactions between tumor cells and CAFs were inferred using CellChat v1.6.1 (https://github.com/sqjin/CellChat) (*34*).

For data mining analyses, single-cell RNA-seq data were obtained from Gene Expression Omnibus (GSE178341). For the analyses shown in Fig. S4G, the data from six pairs of patient-matched tumor and non-tumor samples (three stage III and three stage IV) were were log-normalized by using the NormalizeData() function (Seurat v5.0.3). Data integration was performed using the CCA integration method. Cell type annotation was performed as previously described (*14*). For the analyses shown in Fig. S7E, the data from the tumor samples were used to isolate cancer population, which was further stratified into distinct subpopulations.

### Identification of differentially expressed genes (DEGs)

DEGs for each cell subpopulation were identified by Wilcoxon’s signed-rank test using FindAllMarkers() and shown in table S1, S2, and S4. p-value (< 0.05) and a log2 fold change (log2FC) (> 0.25) were used as cut-off values to select DEGs.

### Gene Set Enrichment analysis

Gene Set Enrichment Analysis (GSEA) was performed by GSEA v1.26.0(https://bioconductor.org/packages/release/bioc/html/fgsea.html). Biological terms listed in MSigDB hallmark gene set (https://www.gsea-msigdb.org/gsea/msigdb/) were imported using msigdbr v7.5.1(https://igordot.github.io/msigdbr/).

### Spatial transcriptomics

Spatial transcriptomic analysis was performed using a Visium spatial gene expression kit (10x Genomics, PN-1000184). Freshly isolated tumor tissues were embedded in OCT compound (Tissue-Tek, #4583), snap-frozen in a bath of 2-methylbutane (Sigma-Aldrich, # 320404) placed on dry ice, and stored at -80. The frozen samples were cryosectioned at 10 μm, placed on a Visium spatial gene expression slide, fixed with cold methanol, and stained with Hematoxylin dye (Agilent, S330930-2). Bright-field images of the slides were captured under a BZ-X800 fluorescent microscope (Keyence). The tissues on the slides were then permeabilized for 25 minutes at 37, followed by *in situ* reverse transcription for 45 minutes at 53. The optimal permeabilization conditions were pre-determined using a Visium spatial tissue optimization slide & reagent kit (10x Genomics, PN1000193). The cDNAs generated *in situ* were collected, and the indexed libraries were constructed using a library construction kit (10x Genomics, PN-1000190). The cDNA libraries were sequenced on the HiSeq2500 platform (Illumina). Raw sequence reads were mapped to the human GRCh38 and mouse mm10 reference genomes (version 2020-A, 10x Genomics) using the Space Ranger pipeline v1.1.0 (10x Genomics), and UMI matrices with spot-associated barcodes were generated.

### Analysis of spatial transcriptomics data

The UMI matrices were loaded into the Seurat pipeline and processed as previously reported(*56*). The spot-level gene expression counts were normalized using the SCTransform() function. The normalized spatial transcriptomics data were processed using the RunPCA() function and subjected to clustering of the spots using the FindNeighbors() and FindClusters() functions with the following parameters: CRC-6_T1, dims = 1:30, resolution = 0.8; CRC-6_T2, dims = 1:30, resolution = 0.74. The processed data were projected in UMAP spaces using the RunUMAP() function. Gene expression levels were displayed in violin plots using the VlnPlot() function or represented spatially on the tissue images using the SpatialFeaturePlot() function. Integration of single-cell RNA-seq and spatial transcriptomics data were performed by using the FindTransferAnchors() and TransferData() functions according to a vignette provided by developers of Seurat software (https://satijalab.org/seurat/articles/spatial_vignette). Pearson’s correlation coefficients to assess the relationship between the prediction scores of the indicated pairs of spatial clusters were calculated using the R package psych v2.4.3 (https://CRAN.R-project.org/package=psych), and the results were visualized using corrplot v0.92 (https://github.com/taiyun/corrplot).

### Analysis of ChIP-seq data

Publicly available ChIP-seq data were obtained from GEO database (https://www.ncbi.nlm.nih.gov/gds/) The following data were re-analyzed by standard protocols: SMAD3 ChIP-seq in HPEK (human primary epidermal keratinocytes) treated with TGF-b3, GSE180252 (GSM5456669, GSM5456670); SMAD3 ChIP-seq in NCI-H441 treated with TGF-b3, GSE51510 (GSM1246712, GSM1246713); SMAD3 ChIP-seq in A549 treated with TGF-b3, GSE51510 (GSM1246720, GSM1246721); SMAD2/3 ChIP-seq in MCF10A MII treated with TGF-b3, GSE83788 (GSM2218835, GSM2218837). Both reference genomes and gene models were obtained from Illumina iGenome website (https://support.illumina.com/sequencing/sequencing_software/igenome.html). FASTQ files were subjected to quality control using FASTP (version 0.23.2) and aligned to the human reference genome GRCh38/hg38 with Bowtie2 (version 2.2.5) with the default parameter. Peaks were called using MACS3 (Model based analysis of ChIP-seq) (version 3.0.0a6) with the following parameters: “-g hs --q 0.01”. Bigwig files with 1-bp resolution were generated and scaled to 1× coverage (reads per genome coverage, 1×RPGC) using bamCoverage function of deeptools package (version 3.5.1) with the following parameters: “--binSize 1 --normalizeUsing RPGC --extendReads 200 --ignoreDuplicates -- effectiveGenomeSize 2701495761”. Input tracks were thereafter subtracted by using deeptools2 bigWigCompare with the following parameters: “--binSize 1 --operation subtract”. ChIP-seq tracks showing peaks distribution on the genome were visualized with PyGenomeTracks (version 3.7) from deeptools.

### Immunofluorescence staining

Tumor tissues or organoid cells were fixed overnight at 4°C in 10% formaldehyde, embedded in paraffin, and sliced into 4 μm sections. After deparaffinization, slides were used for H&E staining or immunostaining. After antigen retrieval with 10 mM citrate buffer (pH 6.0), the slides were stained overnight at 4°C with primary antibodies diluted in D-PBS(-) containing 0.1% BSA and washed with TBST (10 mM Tris-HCl, pH 7.4, 150 mM NaCl, and 0.05% Tween 20). The names and specifications of the antibodies are described in the STAR Methods. Subsequently, the slides were stained with fluorophore-conjugated secondary antibodies for 30 min at room temperature, followed by counterstaining of nuclear DNA with 10 μg/mL Hoechst 33342 (Invitrogen, H3570). In some cases, multiplexed staining with antibodies raised in the same species was performed by sequentially performing the tyramide reaction with the Alexa Fluor Tyramide SuperBoost Kit (Invitrogen, B40922), followed by stripping of the antibodies by 10 mM citrate buffer (pH 6.0) in a pressure cooker for 10 min. Bright-field and immunofluorescent images were captured under a fluorescence microscope (Keyence, BZ-X810).

### Immunohistochemical staining

The deparaffinized slides prepared as described above were subjected to antigen retrieval with 10 mM citrate buffer (pH 6.0) and blocking of endogenous peroxidase activity with 3% H_2_O_2_. After staining with primary antibodies, the slides were stained with biotinylated-secondary antibody and subjected to colorimetric signal detection with Vectastain Elite ABC detection kit (Vector Labs, PK-6100) and DAB tablets (Fuji Film, #040-27001). Nuclear counterstaining was performed with Mayer’s hematoxylin solution (Fuji Film, #131-09665). For semi-quantitative evaluation of the staining of KRT17 and L1CAM, a fraction and intensity of the staining cells at tumor-stromal boundaries of tumors were measured in 10 high power fields.

### Lentiviral transduction

Packaging of pCDH-*luc2*-T2A-BFP (32816913) into lentivirus was performed in HEK293T cells using the ViraPower lentiviral packaging system (Invitrogen, K497500) according to the manufacturer’s protocol (28467911). To establish cancer spheroid cells expressing firefly luciferase and tag-BFP (CRC-6/luc2-BFP or CRC-32/luc2-BFP), the spheroids (CRC-6 or CRC-32) were dissociated using Accumax treatment and infected with lentivirus expressing luc2-T2A-BFP. The luciferase-expressing cells were FACS-sorted based on BFP fluorescence.

### CRISPR/Cas9-mediated gene knockout of cancer spheroid cells

The sgRNA sequences KRT17_#1 (5’-CGCACCTCCTGCCGGCTGTC-3’) and KRT17_#2 (5’-TGCCGGCTGTCTGGCGGCCT-3’), located in the first coding exon of KRT17, were cloned into pX330-U6-Chimeric_BB-CBh-hSpCas9 (Addgene, #42230) at the *BbsI*-digested site to generate pX330-KRT17_#1 and pX330-KRT17_#2, respectively. The sgRNA sequences were designed using the ATUM gRNA design tool (https://www.atum.bio/eCommerce/cas9/input). Control sgRNA vectors were generated by cloning a non-targeting scrambled sequence (5’-GCTTAGTTACGCGTGGACGA-3’) into pX330-U6-Chimeric_BB-CBh-hSpCas9 to create pX330-SC. For targeted disruption of *KRT17* in the spheroids, 1×10^5^ Accutase-dissociated CRC-6/luc2-BFP (32816913) or CRC-32/luc2-BFP cells were transfected with 1 μg of pX330-sg-KRT17_#1, pX330-sg-KRT17_#2, or pX330-SC along with 0.1 μg of pEGFP-N3 (Clontech, PT3054-5) using a NEON electroporation system (Invitrogen). The transfected cells were maintained in 500 μL of spheroid medium for three days, and EGFP-positive cells were single-cell sorted with a FACS AriaIIIu and plated into 96-well plates containing 100 μl of spheroid medium. Aliquots of expanded clones were cultured for 24 h in the presence of 3 ng/mL TGF-β1 (BioLegend, #763104) to examine KRT17 expression by immunoblotting. KRT17-knockout cells (CRC-6_ K17_#1, CRC-6_K17_#2, CRC-32_ K17_#1, CRC-32_K17_#2,) or control cells (CRC-6_Control, CRC-32_Control) were selected.

### Liver metastasis of cancer spheroid cells

The spleens of female NOG mice were injected with CRC-6_Control, CRC-6_K17_#1, or CRC-6_K17_#2 spheroid cells to generate liver metastasis. Briefly, 1×10^5^ cells suspended in 50 μL of a 1:1 mixture of Matrigel and D-PBS(-) were injected into the spleen using a 29 G needle. At 6 weeks post-splenic transplantation, the mice were injected intraperitoneally with D-luciferin (Fuji Film, #120-05114) at 150 μg/g body weight, and growth of liver-metastasized tumors was monitored *in vivo* using an IVIS spectrum imaging system (Caliper Life Science). The data were processed using Living Image software v4.2 (Caliper Life Science). Metastasized livers were harvested for visual inspection.

### Imaging data analyses

Eight-bit grayscale images for each channel were obtained using a fluorescence microscope (Keyence, BZ-X810), and imported into ImageJ (v1.54f), merged into multichannel stacks, and exported as uncompressed OME-TIFF files. These stacks were then imported into QuPath (v0.6.0) (*57*) for automated cell segmentation using the InstanSeg algorithm (v0.1.0) (https://github.com/instanseg). Cell annotation was performed in QuPath using the Single Measurement Classifier, applying marker-specific intensity thresholds to classify cells as positive or negative. Classified cell objects, along with their XY centroid coordinates, were exported as CSV tables. Spatial analyses were conducted in R (v4.2.2), where minimum Euclidean distances from KRT17-positive cells to each marker-positive population were calculated using the sf package (v1.0.21). To quantify infiltration of invasive cancer cells within Vimentin-positive regions, the number of L1CAM TUBB3 cells was normalized to total number of Vimentin cells. All graphical summaries were generated with ggplot2 (v3.5.2).

### Statistical analyses

Data are represented as the mean ± standard deviation. For comparison between two groups, two-tailed Student’s *t* test was performed to calculate *p* values and to determine statistically significant differences. For correlation analyses between the prediction scores shown in Fig. 3G and fig. S3F, Pearson’s correlation coefficients were calculated.

