## supplementary files for "Senolytic targeting of CAF-induced KRT17^+^ colon cancer cells inhibits metastatic invasion"

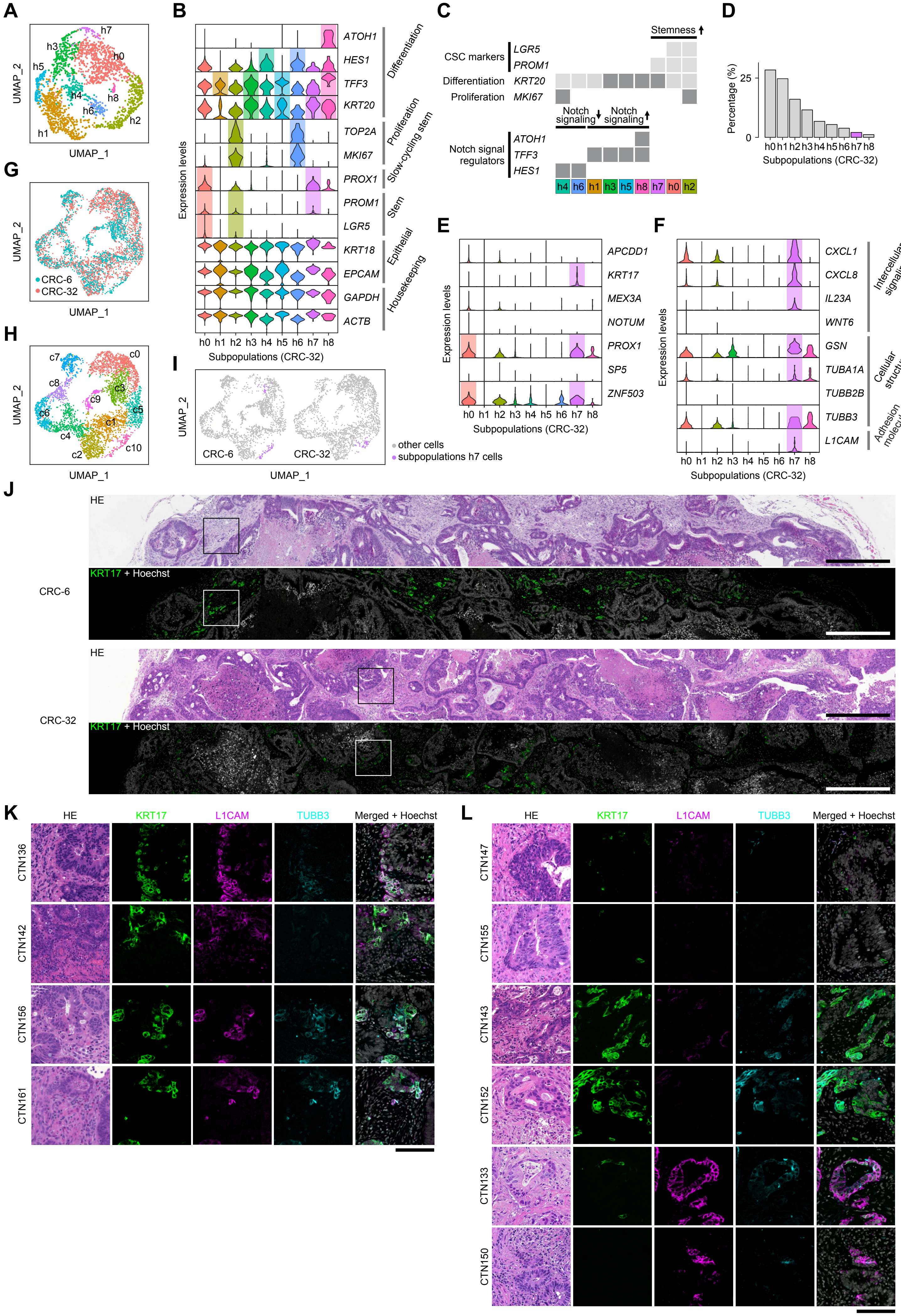


Fig. S1. Shiokawa D. et al.

**Fig. S1. A subpopulation of *KRT17*+ slow-cycling CSCs localizes at the tumor-stromal boundary.**

(**A**) A UMAP plot of the *EPCAM*+ cancer cell subpopulations (h0–h8) of xenograft tumors (CRC-32). (**B**) Violin plots showing expression of representative cell type-specific marker genes in the cancer subpopulations. (**C**) Cell type annotation based on the expression of the cell type-specific markers. (**D**) The relative population of each *EPCAM*+ subpopulation of CRC-32 is shown. The h7 subpopulation is highlighted in magenta. (**E**) Violin plots showing expression of slow-cycling CSC signature genes in tumor cell clusters. (**F**) Violin plots showing expression of genes preferentially expressed in the h7 subpopulation of CRC-6. (**G**) A UMAP plot of the tumor cells from CRC-6 and CRC-32 after data integration. (**H**) A UMAP plot of the cancer cell subpopulations (c0-c10) after the integration of single-cell data from CRC-6 and CRC-32. (**I**) UMAP presentation of the h7 subpopulation of CRC-6 and CRC-32. (**J**) H&E staining and KRT17 immunostaining of xenograft tumors (CRC-6 and CRC-32). The tumor-stromal boundary areas depicted in black (H&E staining) and white (immunostaining) boxes correspond to area shown in Fig. 1H. Scale bar: 500 µm. (**K, L**) H&E staining and immunostaining of surgical specimens of human colon cancers. Staining was performed with the indicated antibodies. Scale bar: 100 µm.

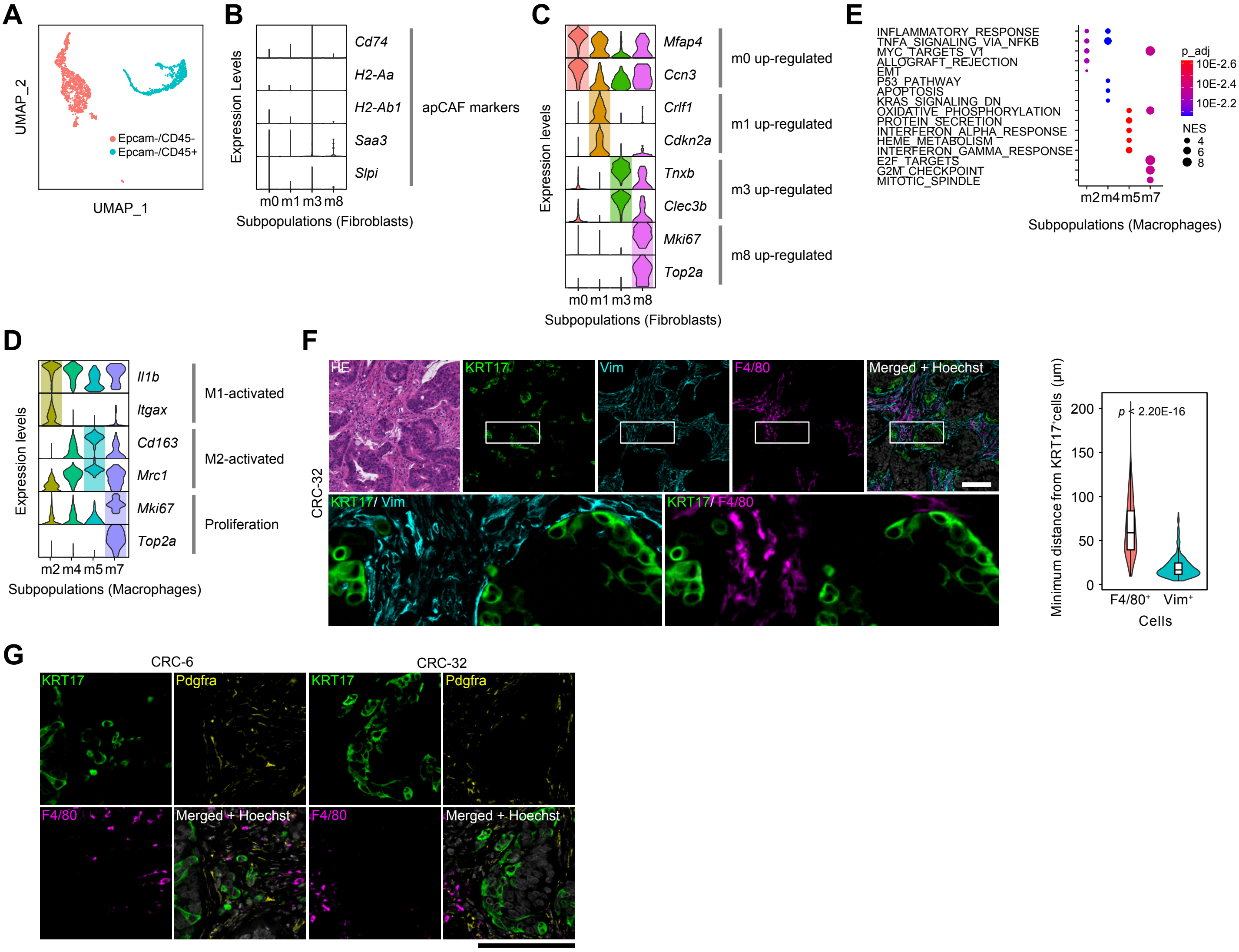


Fig. S2. Shiokawa D. et al.

**Fig. S2. Stratification of stromal populations of the colon tumors.**

(**A**) A UMAP plot showing the distributions of $EpCAM^-/CD45^-$ and $EpCAM^-/CD45^+$ cells (CRC-6). (**B**) Violin plots showing expression of apCAF marker genes in the CAF subpopulations. (**C**) Violin plots showing expression of genes preferentially up-regulated in the CAF subpopulations. (**D**) Violin plots showing expression of M1/M2-associated genes, as well as proliferation markers, in the macrophage subpopulations. (**E**) GSEA of the macrophage subpopulations. (**F**) H&E staining and immunostaining of xenograft tumors (CRC-32). White boxes in the upper panels indicate the regions corresponding to the magnified area shown at the bottom. Staining was performed with the indicated antibodies. Scale bar, 100 µm. Violin plots of the minimum Euclidean distances from $KRT17^+$ cells to $F4/80^+$ and $Vim^+$ cells are shown in the right panel. (**G**) Immunostaining of CRC-6 and CRC-32 xenograft tumors. Staining was performed with the indicated antibodies. Scale bar, 100 µm.

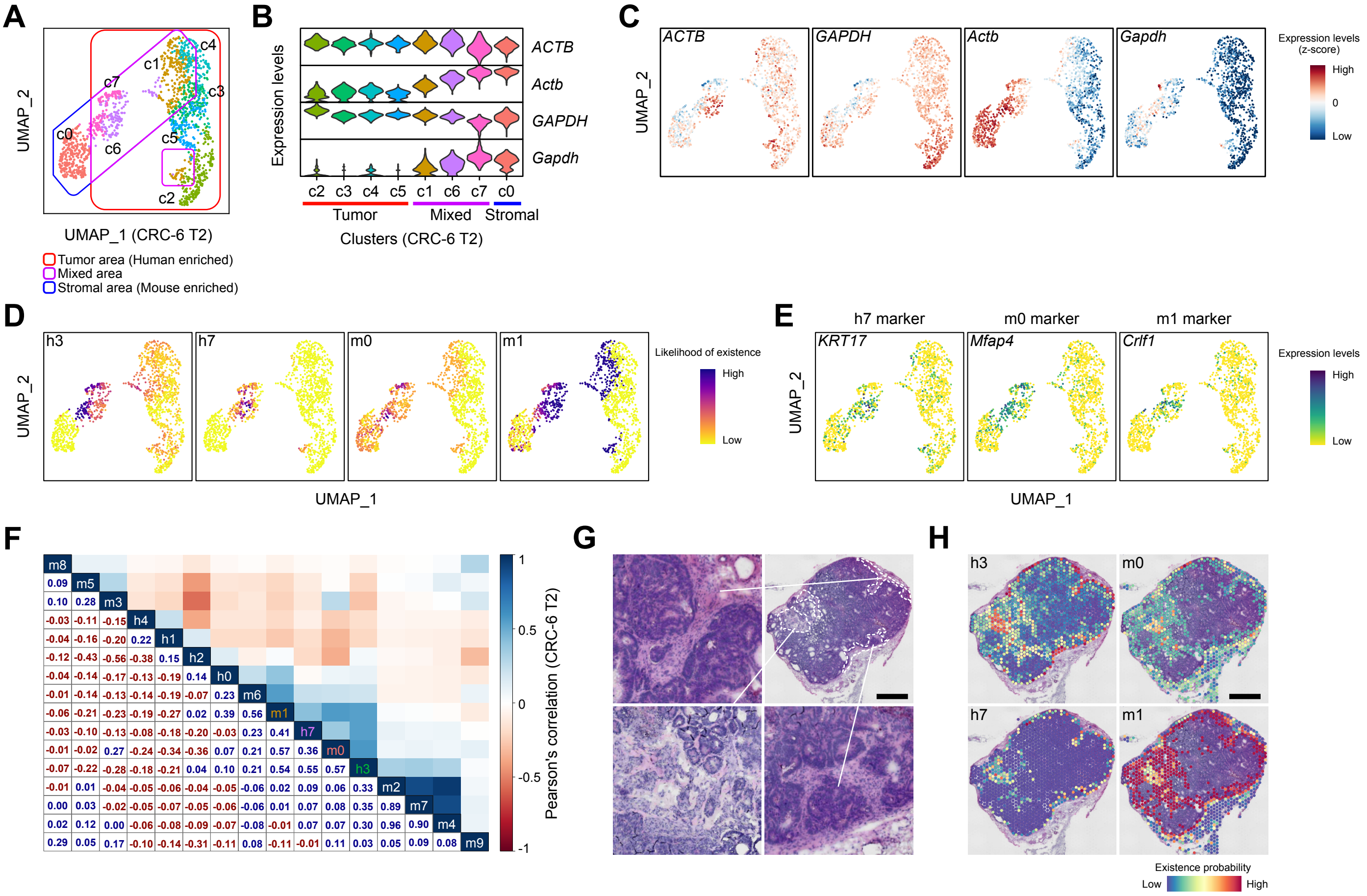


Fig. S3. Shiokawa D. et al.

**Fig. S3. KRT17+ cancer cells colocalize with myCAFs.**

(**A**) A UMAP plot of Visium spots of CRC-6_T2. Eight clusters are denoted by distinctive colors Three types of clusters (a tumor-rich cluster, a stroma-rich cluster, and a mixed cluster) are marked by the indicated brackets. (**B**) Violin plots showing expression of human (*ACTB*, *GAPDH*) and mouse (*Actb*, *Gapdh*) housekeeping genes in the indicated Visium clusters. (**C**) UMAP Feature plots of the Visium spots showing expression of the indicated housekeeping genes (CRC-6_T2). (**D**) Heatmap presentation of the prediction scores for the slow-cycling cancer subpopulations (h3, h7) and the myCAF subpopulations (m0, m1) in UMAP plots of the Visium spots (CRC-6_T2). (**E**) UMAP Feature plots of the Visium spots showing expression of specific markers for the h7 population (*KRT17*), the m0 population (*Mfap4*), and the m1 population (*Crlf1*). (**F**) Correlation matrix showing copresence of the cancer/stromal subpopulations in the Visium spots (CRC-6 T2). Correlation coefficients between the indicated subpopulations are shown as calculated values and color gradients. (**G**) Hematoxylin-stained images of xenograft tumor (CRC-6 T2). Areas surrounded by dashed lines indicate stromal spaces, which are shown in detail in the magnified panels. Scale bar, 1 mm. (**H**) Heatmap presentation of the spatial distribution of the indicated subpopulations in xenograft tumor (CRC-6_T2). Scale bar, 1 mm.

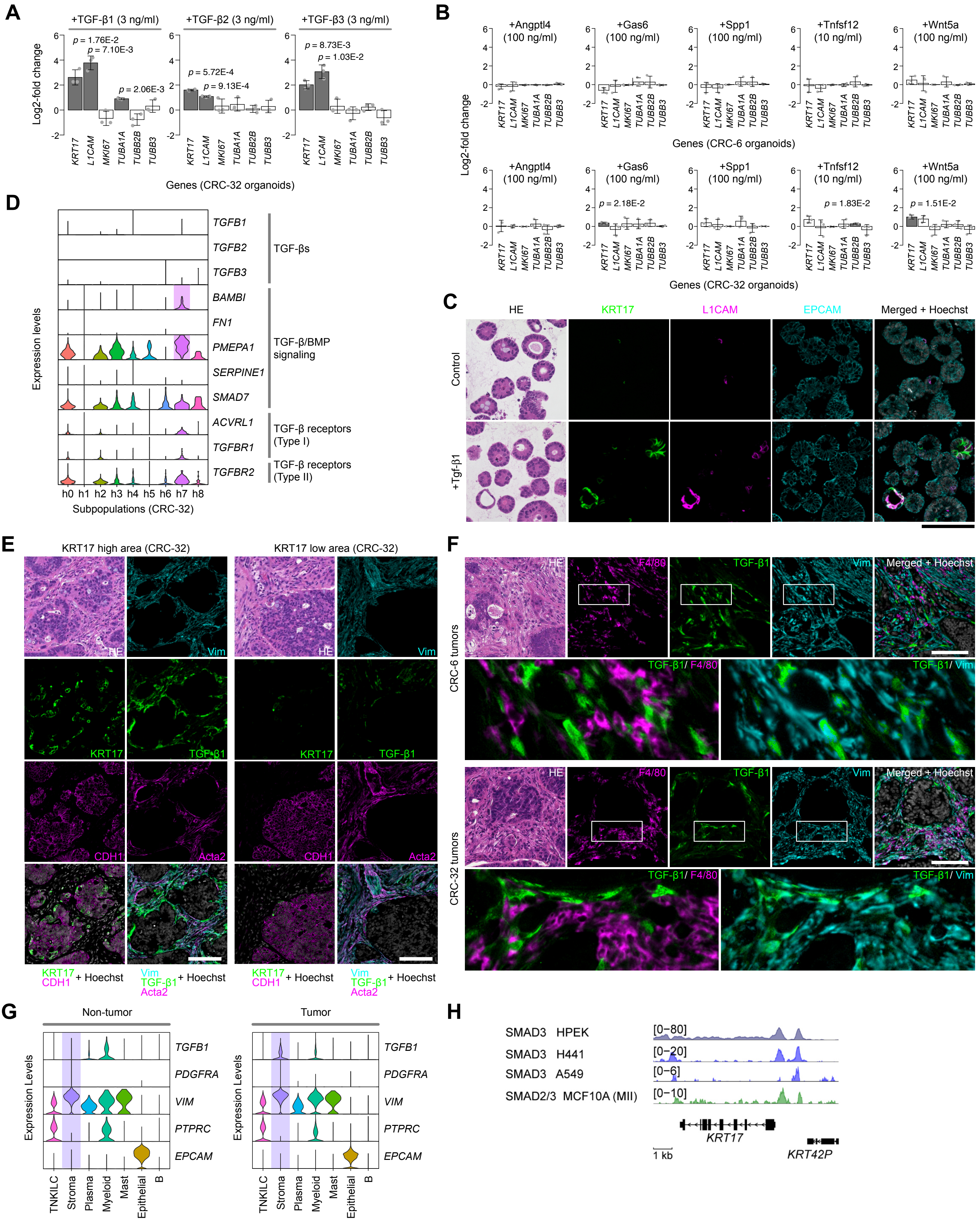


Fig. S4. Shiokawa D. et al.

**Fig. S4. CAF-derived TGF-β1 induces KRT17 expression in cancer cells.**

(**A**) Induction of h7-associated genes by TGF-β. Colon cancer organoids (CRC-32) were treated for 24 h with the indicated isoforms of TGF-β (3 ng/mL) and induction of the indicated h7-associated genes and *MKI67* was evaluated by qPCR (n = 3). Genes induced with statistical significance ($p < 0.05$) are shown as gray bars. (**B**) Induction of the h7 subpopulation-associated genes by myCAF-derived ligands. Colon cancer organoids (CRC-6 and CRC-32) were treated for 24 h with the indicated ligands and induction of the indicated h7-associated genes and *MKI67* was evaluated by qPCR (n = 3). (**C**) Induction of *KRT17* and *L1CAM* in cancer organoids (CRC-32) treated with TGF-β. Organoids were treated for 24 h with vehicle control or TGF-β1 (3 ng/mL) and then used for immunostaining with the indicated antibodies or H&E staining. Scale bar, 100 µm. (**D**) Violin plots showing expression of genes involved in the TGF-β signaling pathway across the cancer subpopulations (CRC-32). (**E**) Colocalization of $KRT17^{+}$ cancer cells with TGF-$\beta1^{+}$ CAFs in colon tumors. Xenograft tumor areas (CRC-6) populated with high or low levels of $KRT17^{+}$ cells were selected for immunostaining with the indicated antibodies or for H&E staining. Scale bar, 100 µm. (**F**) Preferential expression of TGF-β1 in CAFs in xenograft tumors. Tumor/stromal border areas positive for TGF-$\beta1^{+}$ cells were selected for immunostaining and H&E staining. Magnified images of the white boxes are shown in the lower panels. Scale bar: 100 µm. (**G**) Violin plots showing the expression of *TGFB1* and the indicated cell type-specific markers across distinct cell populations from clinical colon cancers. Matched single-cell RNA-seq data (n=7) from tumor tissue (left panel) and nontumor tissue (right panel) of human colon cancers were obtained from public database (GSE178341). The selected single-cell RNA-seq data were then integrated and subjected to cell-type classification into the indicated cell populations. TNKILC collectively represents T cells, NK cells, and innate lymphoid cells. *TGFB2* and *TGFB3* were not detectable in either tissue. (**H**) Binding of SMAD2/3 at the KRT17 gene. ChIP-Atlas database was used to illustrate binding of SMAD2/3 at the KRT17 gene in the indicated human cell lines. Scale bar: 1 kb.

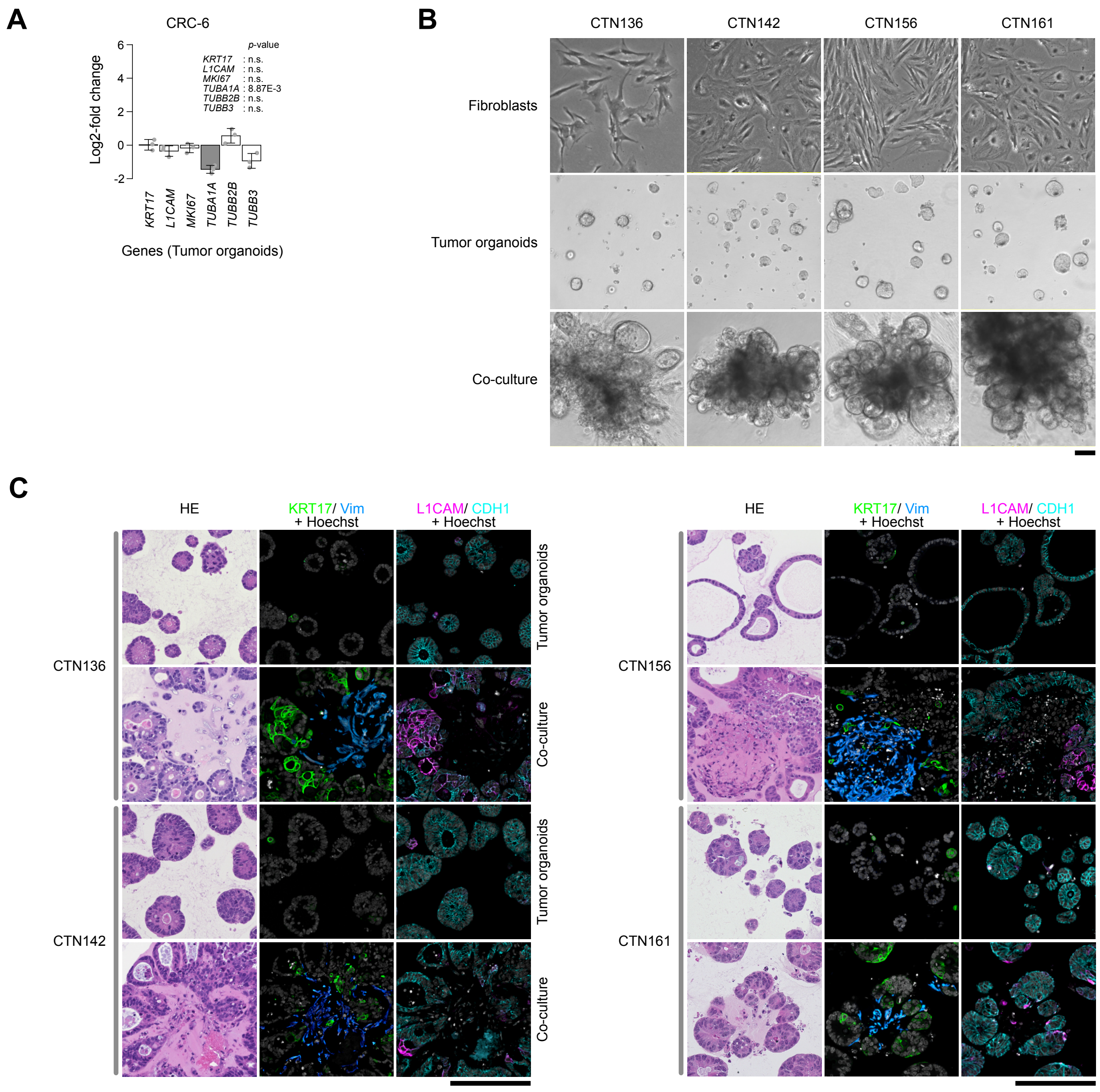


Fig. S5. Shiokawa D. et al.

**Fig. S5. Co-culture with CAFs generates KRT17+ migrating cancer cells.**

(**A**) Effects of coculture with macrophages upon induction of the h7-associated genes in cancer organoids. Cancer organoid cells cultured in the presence or absence of macrophages isolated from xenograft tumors (CRC-6) were used for qPCR for h7-associated genes, and the qPCR values in the absence or presence of the macrophages were compared. Genes induced with statistical significance ($p < 0.05$) are shown as gray bars. n = 3. (**B**) Establishment of *in vitro* coculture of human cancer organoids and CAFs. Colon cancer organoids and CAFs were established simultaneously from surgical specimens (CTN136, CTN142, CTN156, and CTN161). Cancer organoids and matched CAFs were cultivated alone or co-cultivated for 7 days. Phase contrast images of cells are shown. Scale bar: 100 µm. (**C**) Patient-derived cancer organoids grown for 7 days in the presence or absence of human CAFs were used for H&E staining (left panels) or for immunostaining with the indicated antibodies. Scale bar: 100 µm.

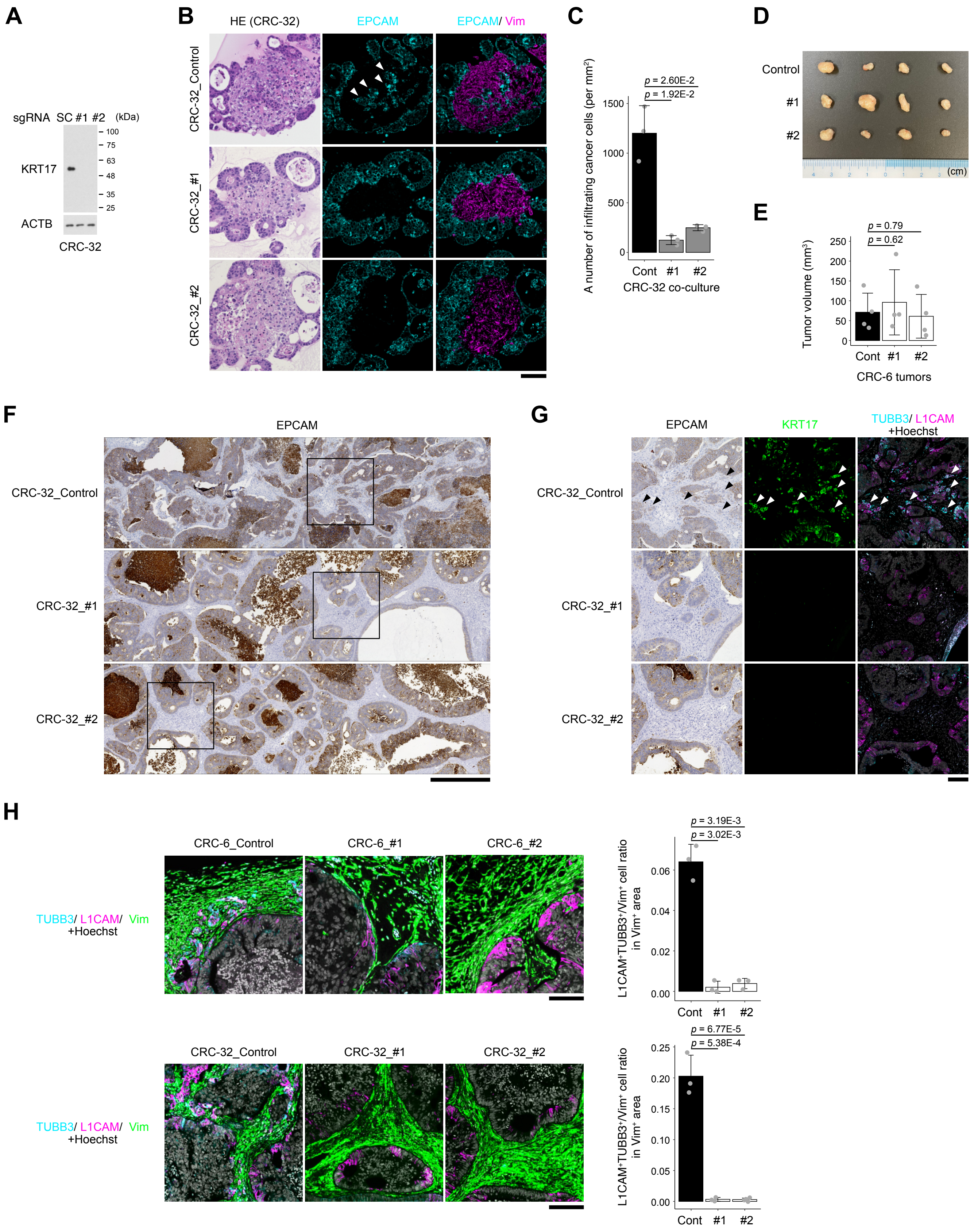


Fig. S6. Shiokawa D. et al.

**Fig. S6. KRT17 is essential for development of invasive cancer cells with metastatic capability.**

(**A**) CRISPR/Cas9-mediated knockout of *KRT17* in cancer cells. Cancer spheroids (CRC-32) expressing Luciferase (CRC-32/luc2-BFP) were introduced with sgRNAs targeting KRT17 (sgKRT17#1, sgKRT17#2) or non-targeting scramble control sgRNA (Control) and then subjected to single-cell FACS sorting and expansion. Cancer spheroids introduced with the indicated sgRNAs were treated with 3 ng/mL TGF-β1 for 24 hrs and subjected to western blot analyses with the indicated antibodies. (**B**) *In vitro* coculture with KRT17-deficient cancer cells. Cancer spheroids described in A were converted into organoids and then used in the co-culture assays with the xenograft tumor-derived CAFs. Formed aggregates were subjected to H&E staining and immunostaining with the indicated antibodies. EpCAM-positive cancer cells infiltrating vimentin-positive stromal areas are indicated by arrowheads. Scale bar: 100 µm. (**C**) The numbers of the infiltrating cancer cells per 1mm$^2$ area shown in B was calculated (n = 3). (**D**) Xenograft tumors generated from control and KRT17-knockout spheroids. NOG mice were transplanted subcutaneously with the indicated CRC-6-derived spheroids. The tumors formed at 3 weeks after transplantation are shown. (**E**) Tumor volumes described in d (n = 4). (**F**) Reduced stromal invasion of KRT17-deficient cancer cells. Xenograft tumors generated from control and KRT17-deficient spheroids described in A were subjected to IHC staining with an anti-EpCAM antibody. Scale bar: 500 µm. (**G**) The boxed areas shown in F were subjected to immunofluorescence staining with the indicated antibodies. Arrowheads indicate invasive tumor cells. Scale bar: 100 µm. (**H**) Immunostaining of control and KRT17-knockout (#1 and #2) xenograft tumors derived from CRC-6 (upper panels) and CRC-32 (lower panels) was performed using the indicated antibodies. Scale bar: 100 µm. The extent of peritumoral invasion by cancer cells in tumor xenografts was evaluated by calculating the relative number of L1CAM$^+$/TUBB3$^+$ tumor cells within the Vim$^+$ CAF$^+$-infiltrated stromal area (the number of L1CAM$^+$/TUBB3$^+$ tumor cells divided by the number of Vim$^+$ CAFs), as shown in the right panels.

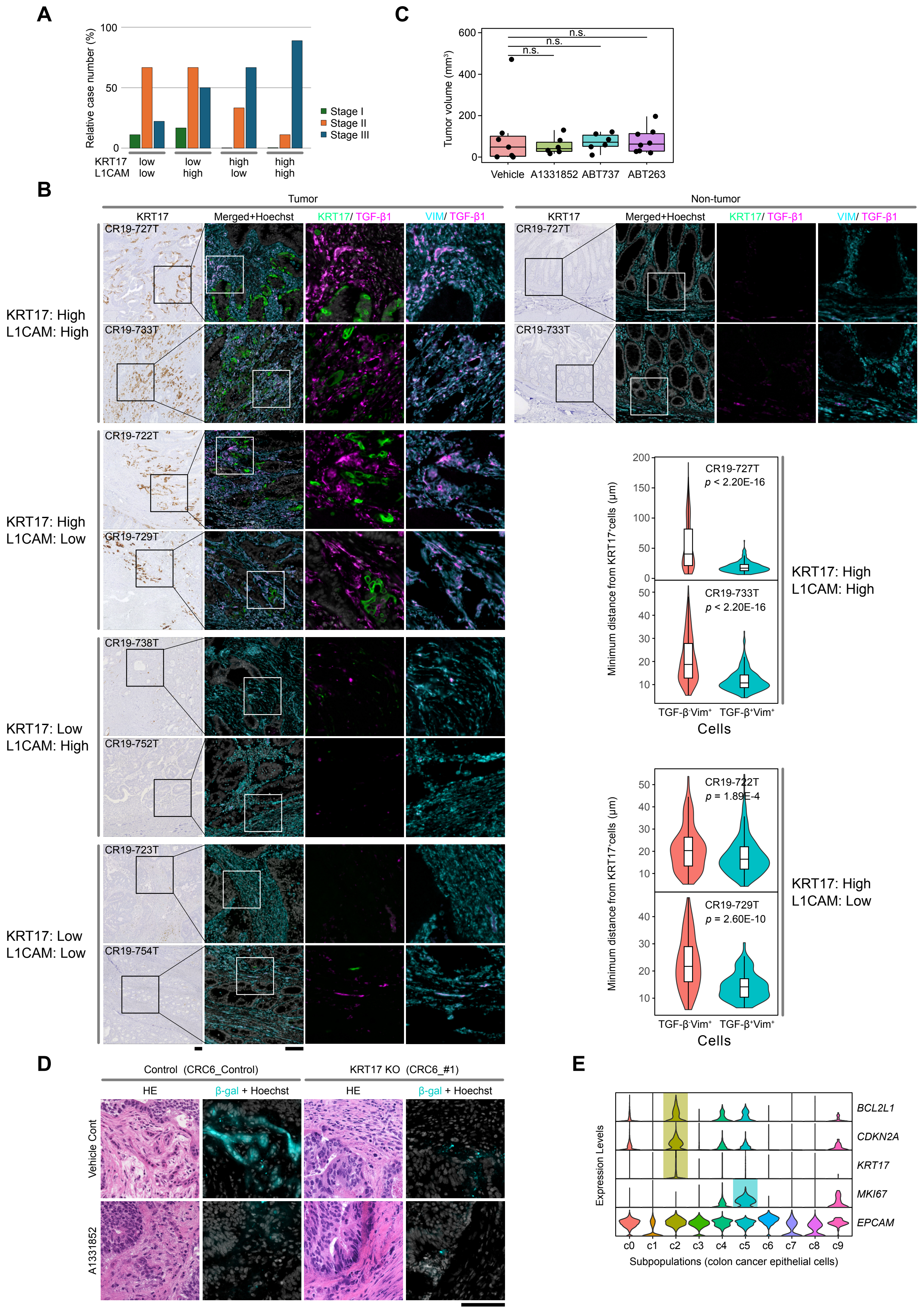


Fig. S7. Shiokawa D. et al.

**Fig. S7. KRT17$^+$ cancer cells colocalize with TGF-β1$^+$ CAFs in clinical colon cancer.**

(**A**) Stage distribution of the four groups described in Fig. 7A. (**B**) Colocalization of TGF-β1$^+$ CAFs with KRT17$^+$ cancer cells in human cancers. The indicated surgical specimens shown in the left column of the IHC staining images (black boxes) were subjected to immunofluorescence analyses with the indicated antibodies. Non-tumor areas of surgical specimens from the KRT17$^{high}$/ L1CAM$^{high}$ cases were also subjected to the IHC and immunofluorescence staining. Magnified images of the while boxes were shown in the right-hand panels. Scale bars: 100 µm. Violin plots of the minimum Euclidean distances from KRT17$^+$ cells to TGF-β$^-$Vim$^+$ and TGF-β$^+$Vim$^+$ cells in KRT17$^{high}$/L1CAM$^{high}$ and KRT17$^{high}$/L1CAM$^{low}$ cases are shown in the right panel. (**C**) Box plots showing the volumes of CRC-6 xenografted tumors treated with the indicated senolytic compounds. (**D**) Fluorogenic staining of SA-β-gal-positive cells in KRT17-deficient tumors. NOG mice were subjected to subcutaneous transplantation of control (CRC6_control) and KRT17-knockout (CRC6_#1) cells for one week, followed by treatment with A1331852 or vehicle control for two weeks before the staining. Scale bar: 100 µm. (**E**) Violin plots showing the expression of the indicated marker genes for KRT17$^+$ senescence-like cells across distinct cancer cell subpopulations from clinical colon cancer samples. Single-cell RNA-seq data from tumor tissues shown in fig. S4G were used to isolate a cancer cell population, which was further stratified into the indicated cell subpopulations.
